# Uncovering features of synapses in primary visual cortex through contrastive representation learning

**DOI:** 10.1101/2022.06.07.495207

**Authors:** Alyssa M. Wilson, Mehrtash Babadi

**Affiliations:** Icahn School of Medicine at Mount Sinai, New York, NY 10029, USA; Work partially performed at Princeton Neuroscience Institute, Princeton University, Princeton, NJ 08540, USA; Broad Institute, Cambridge, MA 02142, USA

**Keywords:** self-supervised contrastive learning, high-throughput electron microscopy, connectomics, mouse visual cortex, chemical synapses

## Abstract

3D EM connectomics image volumes are now surpassing sizes of 1 mm^3^, and are therefore beginning to contain multiple meaningful spatial scales of brain circuitry simultaneously. However, the sheer density of information in such datasets makes the development of unbiased, scalable machine learning techniques a necessity for extracting novel insights without extremely time-consuming, intensive labor. In this paper, we present SynapseCLR, a self-supervised contrastive representation learning method for 3D electron microscopy (EM) data, and use the method to extract feature representations of synapses from a 3D EM dataset from mouse visual cortex. We show that our representations separate synapses according to both their overall physical appearance and structural annotations of known functional importance. We further demonstrate the utility of our methodology for several valuable downstream tasks for the growing field of 3D EM connectomics. These include one-shot identification of defective synapse segmentations, dataset-wide similarity-based querying, and accurate imputation of annotations for unlabeled synapses, using only manual annotation of 0.2% of synapses in the dataset. In particular, we show that excitatory vs. inhibitory neuronal cell types can be assigned to individual synapses and highly truncated neurites with accuracy exceeding 99.8%, making this population accessible to connectomics analysis. Finally, we present a data-driven and unsupervised study of the manifold of synaptic structural variation, revealing its intrinsic axes of variation and showing that synapse structure is also strongly correlated with inhibitory neuronal subtypes.

## INTRODUCTION

Digital reconstructions of brain tissue produced using 3D electron microscopy (3D EM) are proving to be rich resources for learning about properties of neural circuitry^1^. 3D EM labels the lipid membranes and some proteins in biological tissue, producing digital image volumes in which all cells and organelles and some cytoarchitectural elements--including, importantly, components of chemical synapses--are resolved (Fig. 1A)^2,3^. The combined presence of these features allows for highly contextualized analysis of cellular-level neuronal connectivity, and for various organisms, this approach has led to new insights into topics like cell-type-specific nervous system connectivity^3–30^; development- or learning-mediated changes in neural circuitry^31–34^; and the fine structure of neuronal and non-neuronal cells^35–40^.

**Figure 1.**
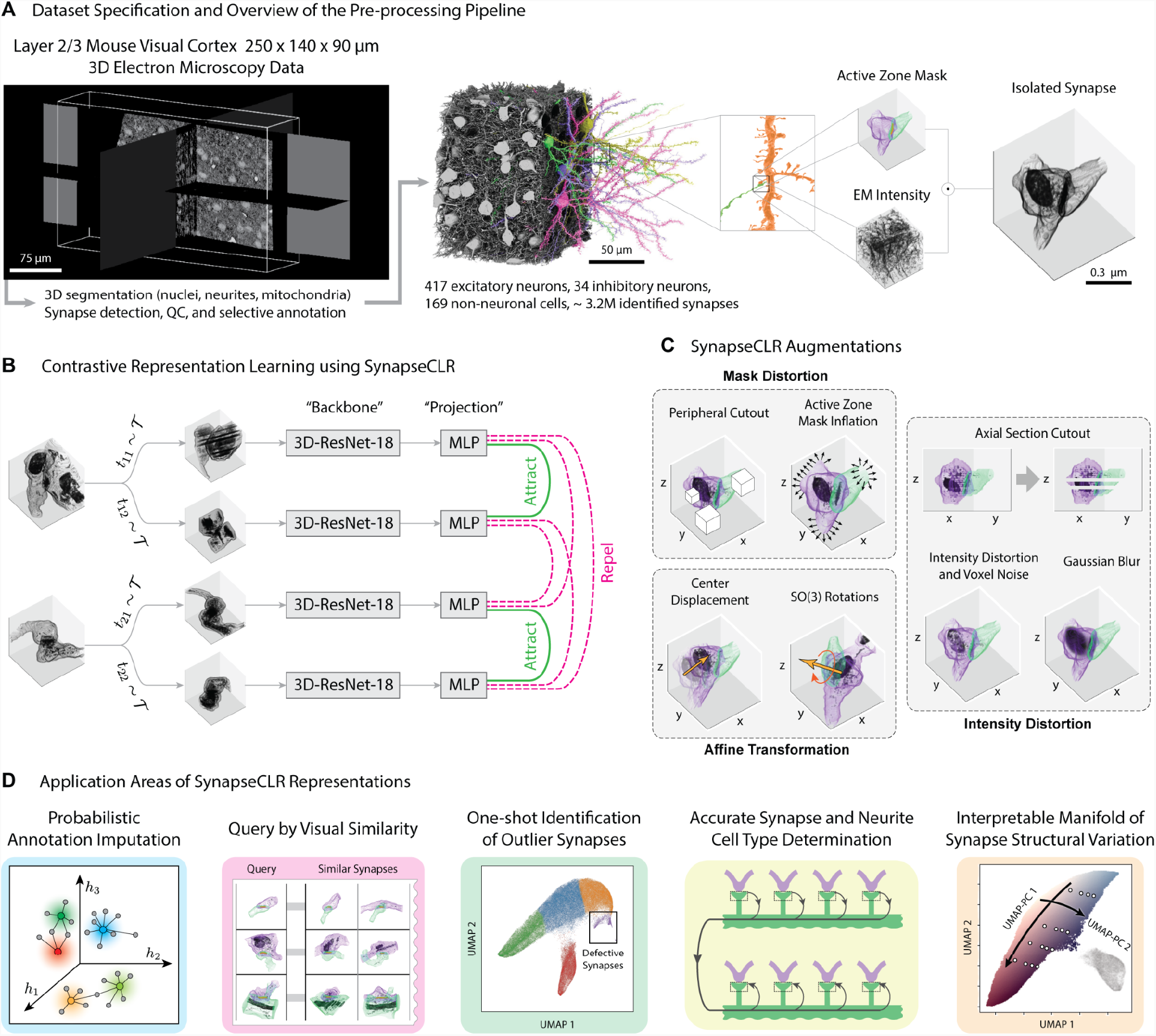
The SynapseCLR workflow. (A) Dataset specification and overview of the pre-processing pipeline; (left) cross-sectional images showing the composition of the 3D EM dataset^2^ used to train SynapseCLR; (middle) a view of this dataset in the same orientation, showing only excitatory neurons (pyramidal cells) with somas in the volume: the left half shows all excitatory somas, and the right half shows 4 examples in color to illustrate the morphologies of these neurons; (middle inset) an example of the synapses in this data from which we drew our training image chunks, in this case involving an excitatory neuron dendrite (orange) that is receiving a synapse from an axon (green); (right) the two raw image chunk layers we used to construct our input image tensors. The top panel shows the segmentation layer, showing only the segmentations of the presynaptic neuron, postsynaptic neuron, and the synaptic active zone (cleft) that they form. The bottom panel shows the EM data from the same region; (far right) we used the segmentation layer to mask the EM, creating an image chunk with the isolated EM image of the synapse only. Our input image tensors were composed of this masked EM chunk. (B) Contrastive learning workflow. Each input image in the tensor is subjected to 2 different, stochastically constructed augmentations (Methods), then passed through a 3D CNN (we used a 3D-ResNet18), and a multi-layer perceptron (MLP) head. The NT-Xent loss used to train the network is built such that the difference between outputs for augmentations of the same image is decreased, and that the difference for augmentations of different images is increased. (C) Augmentations used in SynapseCLR: these include mask distortions, intensity distortions, and affine transformations (Methods). (D) Application areas of SynapseCLR representations explored in the present paper.

Although substantial progress has been made in the annotation and analysis of 3D EM data recently, significant bottlenecks to extracting information from these datasets still exist. For perspective, although most existing datasets span physical scales of only 100s of μm to 1 mm per dimension (1/1000th of a mouse brain and 1/1,000,000th of a human brain)^41^, they are of such high resolutions (∼5-10 nm/px per section) that tera- to petabytes of memory are required to store raw images alone^3,42^. Furthermore, the numbers of biological objects in mid-sized datasets are large: there are typically 10^2^ to 10^4^ cell bodies; 10^6^ “neurites” (i.e. parts of the cells in neural tissue, typically assumed to be pieces of neurons); 10^6^ to 10^8^ of each type of organelle; and 10^6^ to 10^8^ of synapses^2^. These many objects and the many subtle structural variations among objects of each type^7^ make fully human-based analysis of 3D EM datasets infeasible.

For this reason, machine learning approaches for analysis are becoming indispensable in the 3D EM community. Supervised learning approaches have typically been used to detect biological objects in 3D EM data, generating segmentation layers consisting of cells; nuclei; synaptic active zones or clefts; or mitochondria^2,43,44^. To extract neuroscientific insight from these segmentations, however, the main approach has been to measure human-designed features that have been previously identified as structural correlates of function. For example, synaptic cleft size, which is thought to correlate with stronger signal transmission^36,45^, is often a primary measure used to determine the “strength” of connectivity between neurons. Although this approach has led to insights about neuronal connectivity patterns^31,32,35^, it is unclear whether other important structural features are being suboptimally accounted for or missed altogether by relying on human interpretation of past work, which is typically based on highly focused labeling or electrophysiological recordings of particular cells. Furthermore, there is an undetermined correlation or independence between structural measures typically used to characterize synaptic strength such as synaptic cleft size, or the presence of mitochondria^46^.

In this work, we take a step toward building a framework for unbiased, data-driven, and scalable analysis of structural neuronal connectivity patterns using recently developed self-supervised learning (SSL) techniques. Our primary goal here is to learn low-dimensional feature representations of chemical synapses (a fundamental component of neural circuitry) without human supervision, and to evaluate the utility of the obtained representations for key tasks such as identification of parent cell types and prediction of previously noted anatomical measurements. Rather than manually identifying presumably-important and handcrafted features for downstream tasks, or using manual labels to drive the feature learning task, we instead define a collection of structural transformations under which synapse identity should remain “invariant”. Our approach follows the SimCLR framework^47^ that was originally introduced for learning representations of natural images. For this type of task, we employ a convolutional neural network (CNN), which is a flexible artificial neural network for visual feature extraction. We train our CNN to remove redundant visual features while retaining the most robust discriminating visual features using a mix-and-match pretext task (Fig. 1B). Specifically, each input image, in a large batch of randomly selected images, is transformed (“augmented”) twice; all images are then passed forward through the CNN (the “backbone”), followed by an additional shallow multi-layer perceptron (MLP) (the “projector”) for further feature processing, and the resulting representations are used to evaluate and minimize a contrastive loss function. The loss function is designed such that *similarity* between representations from the *same* source image, and *dissimilarity* between representations from *different* source images, both lead to lower loss (Methods). In practice, SSL frameworks have been shown to produce powerful low-dimensional visual representations that match and even surpass those obtained using other machine learning paradigms, such as supervised learning^48^. The SSL representations have been shown to be applicable for a wide range of visual tasks such as classification and segmentation^47,49–52^ and few-shot transfer learning^53,54^. The success of the SSL paradigm largely relies on judicious choice of augmentation transformations (e.g. rotation, random cropping, or color histogram distortions), which can be highly domain-specific, as well as on the implicit inductive bias of the backbone artificial neural network. A natural artificial neural network architecture for visual data processing is the CNN, owing to its desirable translational equivariance and hierarchical feature extraction properties.

We adapted the SimCLR framework to work on 3D volumetric data rather than 2D images, along with a set of augmentation transformations specifically designed for segmented 3D EM data (Fig. 1C; Methods). We used the resulting package, called “SynapseCLR”, to train a 3D-ResNet18 CNN^55^ on 97658 synapses selected from a recently released 3D EM reconstruction of mouse primary visual cortex^2^. These representations reveal that synapses are partitioned into several highly distinct regions in the feature space, with smooth variation of representations within each region (see Results). Importantly, by comparing these representations with annotated anatomical measurements for 5664 of these synapses that were verified previously^2^, we find that the structures that synapses occupy in feature space correspond to changes in structural correlates of function. Excitatory-to-excitatory synapses; synapses involving inhibitory neurons; and synapses with segmentations contaminated by breaks or other artifacts each occupy distinct regions of the representation space. Within each connected component of the representational manifold, we observe smooth and correlated changes in synapse size, the presence of pre- and postsynaptic mitochondria, and the distance between a synapse and its postsynaptic neuron. This emergent low-dimensional manifold structure allows us to map out relationships between the various structural annotations we have at our disposal for this study. The strong correlation between these annotations and the representations allows us to use them for a variety of downstream tasks (Fig. 1D), including one-shot detection of synapses with defective 3D reconstructions in a dataset; imputation of structural measures for unlabeled synapses by regressing against available annotations; accurate cell type assignment for small neurites that form synapses but otherwise lack the morphological context needed for typing--this application allows those neurites to be included in neural connectivity analysis; and querying of synapses across a dataset based on their visual similarity.

The SynapseCLR package is publicly available at https://github.com/broadinstitute/SynapseCLR, and can be readily used for learning representations from other 3D EM datasets.

## RESULTS

We pulled synapse images from an EM dataset^2^ consisting of ∼2250 40-nm-thick tissue sections (resolution 3.58 × 3.58 nm^2^/px^2^) stacked into a total volume of 240 × 140 × 90 μm^3^ (Fig. 1A). This dataset has multiple segmentation layers, three of which we used: first, a cellular segmentation containing 451 neurons with somas in the volume and ∼8 million neurites; second, a segmentation of ∼3.2 million automatically detected synaptic clefts; and third, a segmentation containing ∼2 million automatically detected mitochondria. Of the synaptic clefts, 5664 had been previously verified by experts as representing real synapses, because they were formed between neurons with somas in the volume (398 total) and were used for previous analysis of connectivity between those cells. Biological annotations had also been made and verified for these synapses, including pre- and postsynaptic cell type (i.e. excitatory vs. inhibitory); distance of the synapse from the pre- and postsynaptic somas; synaptic cleft size; and number and size of pre- and postsynaptic mitochondria, i.e. mitochondria located within ∼1-5 μm of the synaptic cleft.

To maximize our ability to evaluate and interpret our learned representations, we included all 5664 of these synapses (called “annotated synapses”) in our training, along with 91994 additional synapses we randomly selected from the remaining clefts (“unannotated synapses”; Methods). These ∼10^4^ unannotated examples were located throughout the volume, thus providing a spatially uniform sampling of the synapses across different cell types and different regions within this piece of cortex (layer 2/3 of area V1).

### SynapseCLR learns meaningful low-dimensional representations of synapses

We structured each input image beginning from a two-channel 3D image tensor centered at the synaptic cleft centroid. The first channel was an integer encoding of the presynaptic process, synaptic cleft, and postsynaptic process segmentation layers (Fig. 1A). The second channel was the volumetric EM image chunk from the same region. After augmentation and preprocessing, each image that was ultimately used for training was a 3D image chunk containing only the EM of the presynaptic process, synaptic cleft, and postsynaptic process (Fig. 1A,C, Methods). We masked our EM images in this way because we wanted to learn representations of chemical synapses in isolation. Although the dense neuropil that surrounds each synapse may impact the physiology of that synapse to varying degrees--microglia can modulate synaptic function^56^, whereas many axons pass closely to synapses without forming any apparent functional interfaces^5,57^--the entropy of this external material is so high that it would be challenging to learn a meaningful representation of the synapse itself if it were included. Studying potential interactions between synapses and the surrounding processes can be done as a secondary task, which requires having meaningful representations of isolated synapses in the first place (the focus of the present work).

We developed a set of augmentation transformations specific to synapses that included both spatial/geometric and intensity transformations, since combinations of both types lead to the most effective representations^47^ (Fig. 1C; Methods). Each augmentation was a stochastic composition of global and per-section brightness adjustment; additive and multiplicative pixel-level noise; Gaussian blur; corner and shell masking cutouts in order to encourage learning part-whole relationships; dilations of the segmentation mask applied to the EM image channel; single-section masking; affine transformations (translations and arbitrary 3D rotations, without shearing or scaling since such deformations would mask known identifying properties of synapses); x-axis flip; y-axis flip; and x-y swap. Some of these transformations involve hyperparameters, which were stochastically selected from ranges chosen within reason for each augmentation (Methods).

We trained the 3D-ResNet18 backbone and the 2-layer MLP projection head (see Fig. S1A) on all 97658 synapse image tensors across 4 NVIDIA Ampere A100 GPUs, each with 96GB GPU RAM. We used a batch size of 192 and trained the network for 200 epochs over two weeks, with a learning rate restart after 100 epochs. The loss function decreased steadily and reached a stable minimum (Fig. S1B; Methods).

Following successful training, one can take activations from any layer of the SynapseCLR network as a potentially useful representation, with the general expectation that deeper layers contain further processed and aggregated information. Here, we explored the 512-dimensional feature maps from the last layer of the 3D-ResNet18 backbone, as well as the 512-dimensional middle layer, and the 128-dimensional last layer of the projection head. In each case, we generated UMAP^58^ embeddings from all representations, and found that the embeddings were structurally consistent (Fig. S2). Further quantitative benchmarks revealed that the 512-dimensional backbone features provided the highest utility for predicting biological properties of unannotated synapses (see below). As such, we primarily base our main discussion on backbone features, and briefly comment on the utility of other features later (Discussion). We explored both 2D (Fig. 2A) and 3D (Fig. S3) UMAP embeddings of these representations and found in both cases, these representations occupied several highly separable regions in feature space. A detailed analysis of synapses in each region will be provided later; examples from each region are shown in Fig. 2B-C.

**Figure 2.**
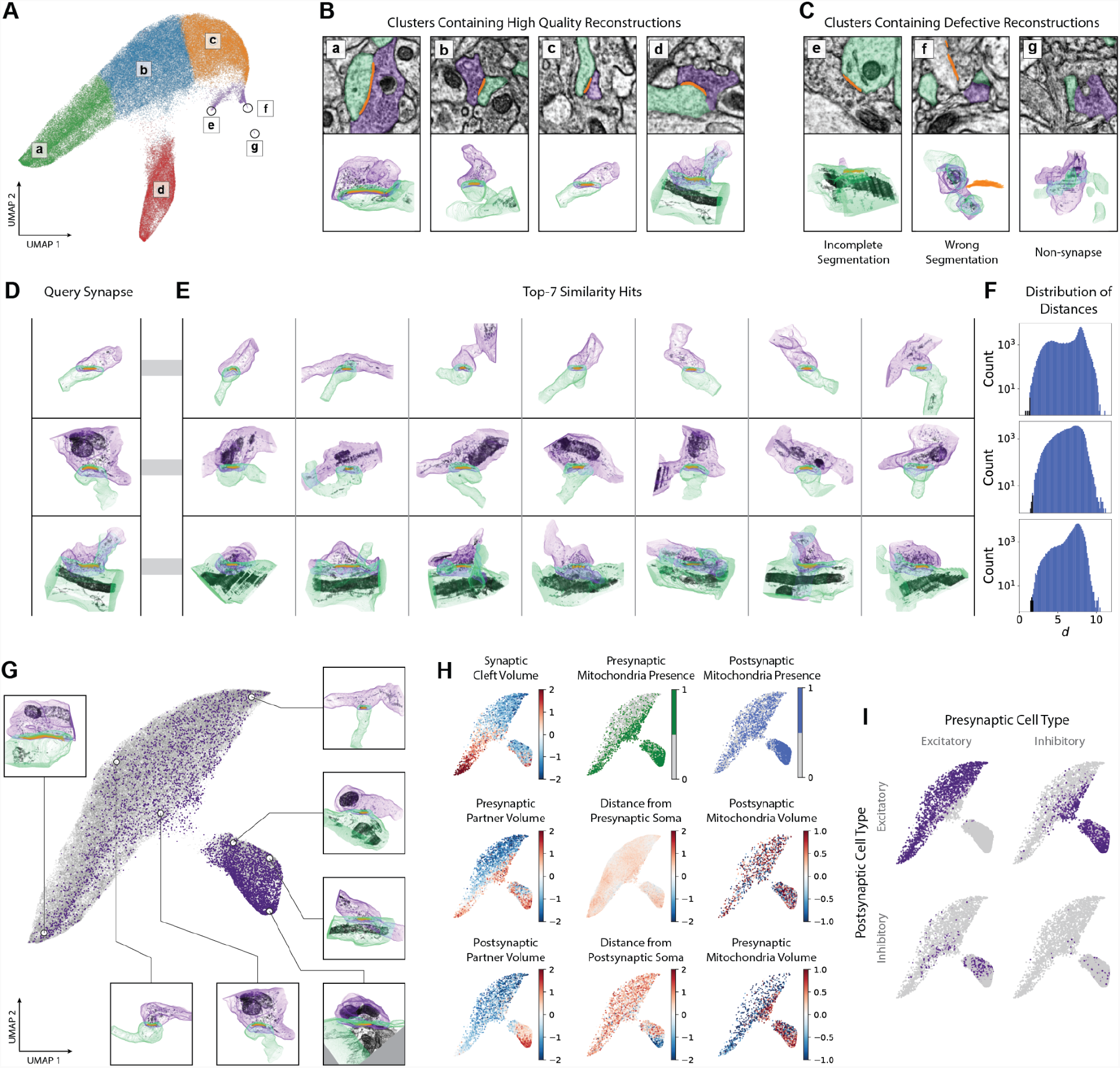
Relationships between SynapseCLR representations and structural properties. (A) 2D UMAP embedding of representations obtained from the trained 3D-ResNet18 backbone features, clustered by building a *k*-nearest-neighbor graph (k=100) according to cosine similarity and clustering using the Leiden algorithm. Circles around e, f, and g are guides for the eye. (B) Example images of synapses from clusters that contain primarily high-quality reconstructions (a-d). (C) Example images of synapses from clusters that contain defective reconstructions (e-g). (D) Segmentation-masked EM for 3 arbitrary synapses used as “prompts” to query structurally similar synapses in the representation space. (E) The 7 nearest neighbors in representation space for each prompt synapse in (D), showing remarkable visual similarities to the corresponding prompt synapse. (F) Representational Euclidean distances between each synapse and the corresponding prompt synapse shown in (D). Distances for the 7 nearest neighbors shown in (E) are colored black in the histogram. (G) 2D UMAP of representations for only non-defective synapses (94874 in total). Purple points: the 5623 high-quality synapses for which we had annotations. Gray points: the 89251 high-quality synapses without annotations. (H) Each structural property measured for annotated synapses, superimposed on the 2D UMAP. Points representing the locations of annotated synapses formed between neurons of various valence (dark purple) in the UMAP, showing substantial separation of synapses in representation space as a function of pre- and postsynaptic neuron type.

As a first visual assessment for the meaningfulness of the learned representations, we selected a few typical synapses from the dataset as “prompts”, and “queried” similar synapses based on Euclidean distance in the representation space (Fig. 2D-F, Fig. S4). We noticed that the queried neighbors shared staggeringly similar structural features to the prompt synapses, providing a first demonstration that SynapseCLR representations highlighted features that capture structural variability across synapses in the dataset.

### SynapseCLR representations allow one-shot identification of contaminated synapse segmentations

In our inspection of regions in these UMAP projections, we discovered that one in particular contained primarily synapses with segmentations that were contaminated by artificial breaks or other imperfections (Fig. 2B,C). Crucially, several types of contaminated synapses formed a single cluster in the representation space, allowing us to remove them from the dataset in a single operation, along with 5 nearest neighbors as a conservative quality control measure (Methods). The number of annotated and unannotated synapses passing QC amounted to 5623 and 89251, respectively. The removed contamination amounted to 0.7% of proofread and manually annotated synapses, compared to 3% of the remaining synapses, demonstrating the specificity of this quality control measure. We did not find synapses exhibiting similar issues in other regions of the representation space.

### SynapseCLR representations correlate with meaningful anatomical features

To assess the relationship between these learned representations and anatomical measures that are conventionally used as structural correlates of function, we overlaid each property measured for the 5623 high-quality annotated synapses (Fig. 2G) on our 2D UMAP embedding (Figs. 2H, I).

We found that synapses formed between different neuron types (excitatory/E or inhibitory/I; Methods) occupy largely distinct regions of representation space (Fig. 2I). Specifically, E-to-E synapses dominate the larger, upper manifold, and synapses involving inhibitory neurons (I-to-E, E-to-I, and I-to-I) dominate the lower manifold as well as much of the thin bridge connecting the two. Among the domains inhabited by synapses involving inhibitory neurons, we observe some additional type-specific representation partitioning: I-to-E synapses appear to concentrate in the lower part of the bottom manifold, whereas E-to-I and I-to-I synapses appear to be mostly excluded from this subregion. This representational organization implies that synapses have inherently distinct structures depending upon whether they are formed by an excitatory or inhibitory neuron and upon whether the synaptic target is an excitatory or inhibitory neuron (see Discussion).

Within each of the two largely separate representation manifolds, we also observed structured variation of other annotated measures. Notably, synaptic cleft size varied smoothly along the longest dimension of the upper (E-to-E) manifold. We observed a similar but less dramatic change in cleft size in the primarily-I manifold, again along the longest dimension. The sizes of presynaptic and postsynaptic partners also co-varied somewhat with cleft size in both manifolds (Fig. 2H, left column), but, notably, we see that large presynaptic and postsynaptic interfaces do not always correspond with a large cleft size (e.g. at the bottom of the lower manifold).

We also observed orchestrated variation in whether mitochondria were present in pre- and postsynaptic partners, and in the volume of those mitochondria that was captured in our image chunks (which is a rough proxy measure of how close mitochondria are to the synaptic cleft). In particular, presynaptic mitochondria generally co-occurred with large presynaptic processes in synapses of any type (Fig. 2H top-middle, left-middle).

In contrast, the learned synapse representations captured limited information related to synaptic distance from the presynaptic soma (Fig. 2H, middle). Distance from the postsynaptic soma also varied little except at the bottom of the I manifold, where distances were minimal compared to those for other synapses (Fig. 2H, bottom middle).

To further ascertain the value of our SynapseCLR representation learning strategy, we also obtained baseline representations using two other CNNs: a random (untrained) 3D-ResNet18 CNN, and MedicalNet^59^, a general-purpose pretrained 3D-ResNet18 CNN for 3D segmentation of MRI and CT datasets in any of a diverse set of organs (including brain). Our inclusion of MedicalNet as a baseline is motivated by the success of “transfer learning” approaches, i.e. the empirical observation that CNNs trained on images from one domain, e.g. the ImageNet natural image dataset^60^, exhibit strong generalization to other imaging data domains, such as histopathology slides^61^ and cell morphology images^62^.

The resulting UMAP embeddings of SynapseCLR representations and the two baselines, along with several overlaid annotations, are given in Fig. S2. We notice significantly stronger correlation and structure in SynapseCLR representations compared to the two baselines. A quantitative benchmark of these representations for imputing anatomical annotations will be presented in the next section.

Given that our SynapseCLR representations are robust descriptors of variation amongst synapses in the dataset (by design), we conclude that annotations that vary smoothly across representation space (like synapse size and mitochondrial presence) are indeed important factors in establishing the structural identity of a synapse.

### SynapseCLR representations accurately impute properties of unannotated synapses

The strong correlations we observe between SynapseCLR features and anatomical annotations suggest the possibility of using these features to predict (“impute”) annotations for unannotated synapses (which comprise the vast majority of synapses in our dataset). Endowing synapses with anatomical data is useful for bypassing the need for manual annotation and for increasing the number of synapses that can be used to assess connectivity in the 3D EM dataset. Although our set of annotated synapses is biased (it is composed of only synapses between neurons with somas in the EM volume, i.e. synapses between local neurons), we see that representations of annotated and unannotated synapses occupy all of the same domains and rather uniformly in feature space (see purple and gray points in Fig. 2G). Thus, although other synapses in the volume may be formed by physically more distal cells of differing subtypes, there are no systematic structural features they possess that are not also represented in the annotated synapse population. For this reason, annotated synapses can be used to impute properties for these other synapses.

To generate imputed values of each type of annotation for our unannotated synapses, we trained Gaussian process (GP) regression models using the SynapseCLR representations as covariates and the annotations from our subset of 5623 annotated synapses as targets. We optimally regularized each model, including the choice of GP kernel, using a cross-validation strategy (Methods). For comparison, we also trained regression models using representations from the two other baseline CNNs. We found that the backbone-level representations from the trained SynapseCLR CNN generated the best predictions overall (Fig. 3A; Table S1; Methods). In the best models, the area under the ROC curves (AUC) for predicting pre- and postsynaptic cell types were 0.975 +/- 0.004 and 0.940 +/- 0.004 AUC, respectively (Methods). The presence of pre- and postsynaptic mitochondria could also be predicted with high accuracy (0.939 +/- 0.015 and 0.857 +/- 0.001 AUC, respectively). Another annotation which could be predicted well was synaptic cleft size (70.0 +/- 2.7% explained variance). On the other hand, our models for a synapse’s distance from its pre- and postsynaptic neuron somas explained only 15.9 +/- 1.4% and 53.9 +/- 1.6% of the variance in those measures, respectively. Looking at the scatter plots of predicted vs. actual pre- and postsynaptic soma distances (Fig. 3C), it seems that both regression models are similarly limited in their ability to impute distances except for the subset of synapses that have small distances to the postsynaptic soma (the blue regions in the third column of Fig. 3B). Overall, the diminished predictability of synaptic distances to somas from their structure suggests that the correlation between these two features is subtle. Our models for predicting pre- and postsynaptic mitochondria volumes explained 56.9 +/- 3.1% and 10.0 +/- 2.4% of the variance in those measures, respectively. The limited predictive power of these models likely stems from our use of small synaptic image chunks. We will discuss potential future improvements in the Discussion section.

**Figure 3.**
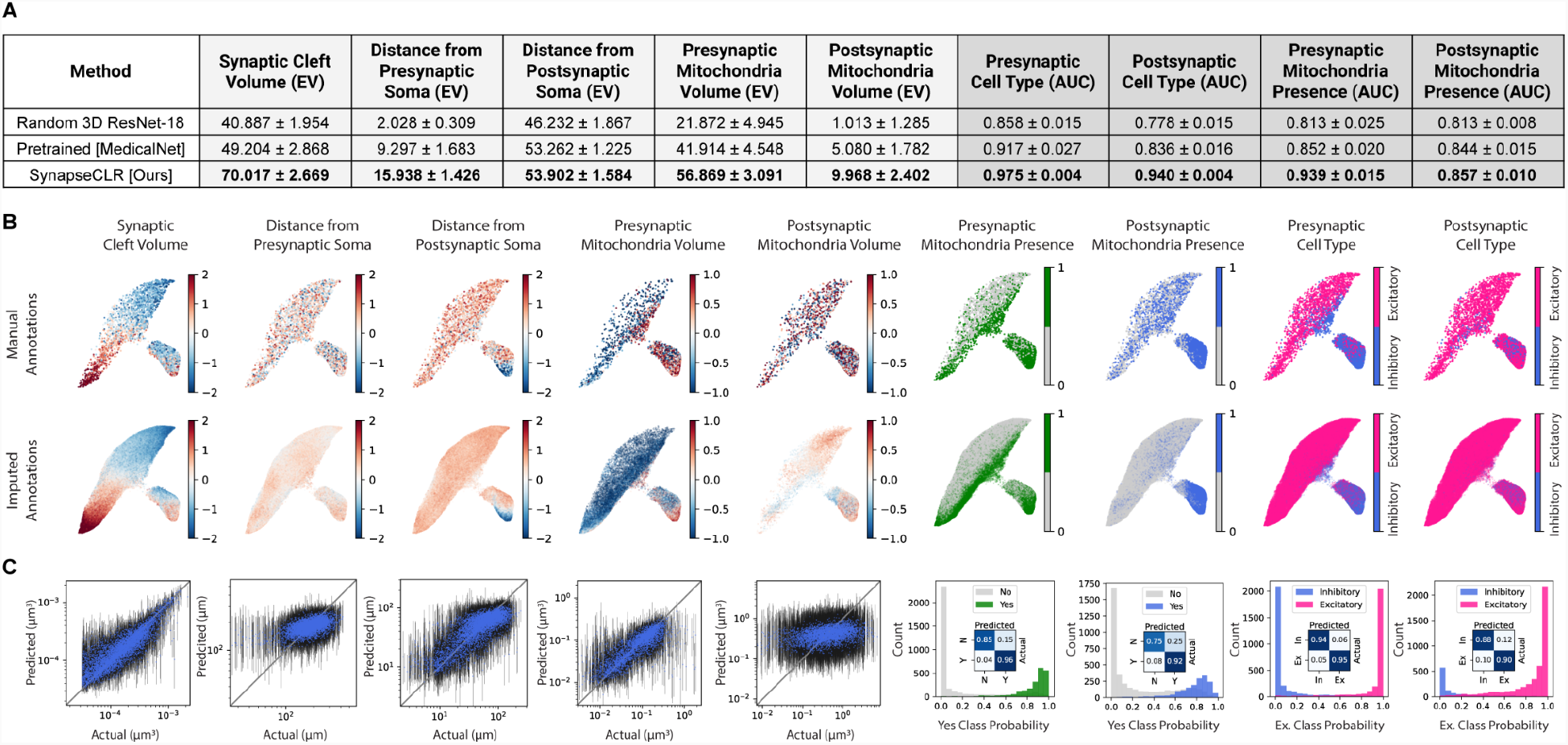
Predicting structural properties of unannotated synapses using SynapseCLR representations as covariates and Gaussian process regression. (A) Annotation prediction accuracy for the best Gaussian process regression models trained on synapse representations using (top row) a random 3D-ResNet18, (middle row) a general-purpose pretrained 3D-ResNet18 CNN for 3D segmentation of MRI and CT datasets (MedicalNet), and (bottom row) SynapseCLR 3D-ResNet18 backbone representations. Light gray columns: explained variance (EV) for continuous measures. Dark gray: area under the receiver operating characteristic curve (AUC) for categorical measures. Bold: the best-accuracy model for each annotation. (B) UMAP embedding of SynapseCLR 3D-ResNet18 backbone representations with structural properties on the subset of annotated synapses (top), and imputed structural properties for all synapses (bottom). (C) Imputed vs. measured structural values for annotated synapses. Blue markers show the Gaussian process posterior mean and the black error bars show the 90% credible intervals. The right-most four panels correspond to binary measures. The histograms show the predicted probabilities stratified by true class labels; the inset plot shows the binary classification confusion matrix.

### Consensus calls of imputed cell types from multiple synapses increases the population of analyzable neurites in 3D EM datasets

Of these imputed annotation values, the ones that are most important for growing a connectomics dataset are the pre- and postsynaptic cell types: E and I neurons are generally understood to have very different functional roles in neuronal circuitry, so any neurite that cannot be typed is excluded from analysis because its role in the circuit cannot be ascertained.

However, this conservative approach leads to a severely reduced picture of neuronal circuitry. Many of the synapses in this EM dataset (all ∼3.2 million of them, besides the 5623 annotated synapses between cells with somas, that is, less than 0.2% of all synapses) are formed between neurites that cannot be assigned a valence from morphology, precisely because their somas are not inside the imaged volume. The result is that >99.8% of synapses in the dataset are not being used to describe circuitry characteristics.

In the previous section, we showed that for individual synapses, SynapseCLR representations can be used to reliably predict pre- and postsynaptic cell types (0.975 and 0.940 AUC, respectively). Here, we exploit the fact that a typical synapse-forming neurite in this dataset that is not connected to a soma contains tens of synapses (for neurites connected to a soma, this figure reaches thousands of synapses; Fig. 4C). Therefore, even with imperfect cell typing for individual synapses, our ability to confidently assign a cell type to each neurite will be dramatically enhanced (with errors decreasing exponentially with the number of synapses per neurite) by taking a “consensus” of all participating synapses in a maximum likelihood estimation fashion. The consensus cell type of each neurite can then be propagated back to the participating synapses (Fig. 4A; Methods).

**Figure 4.**
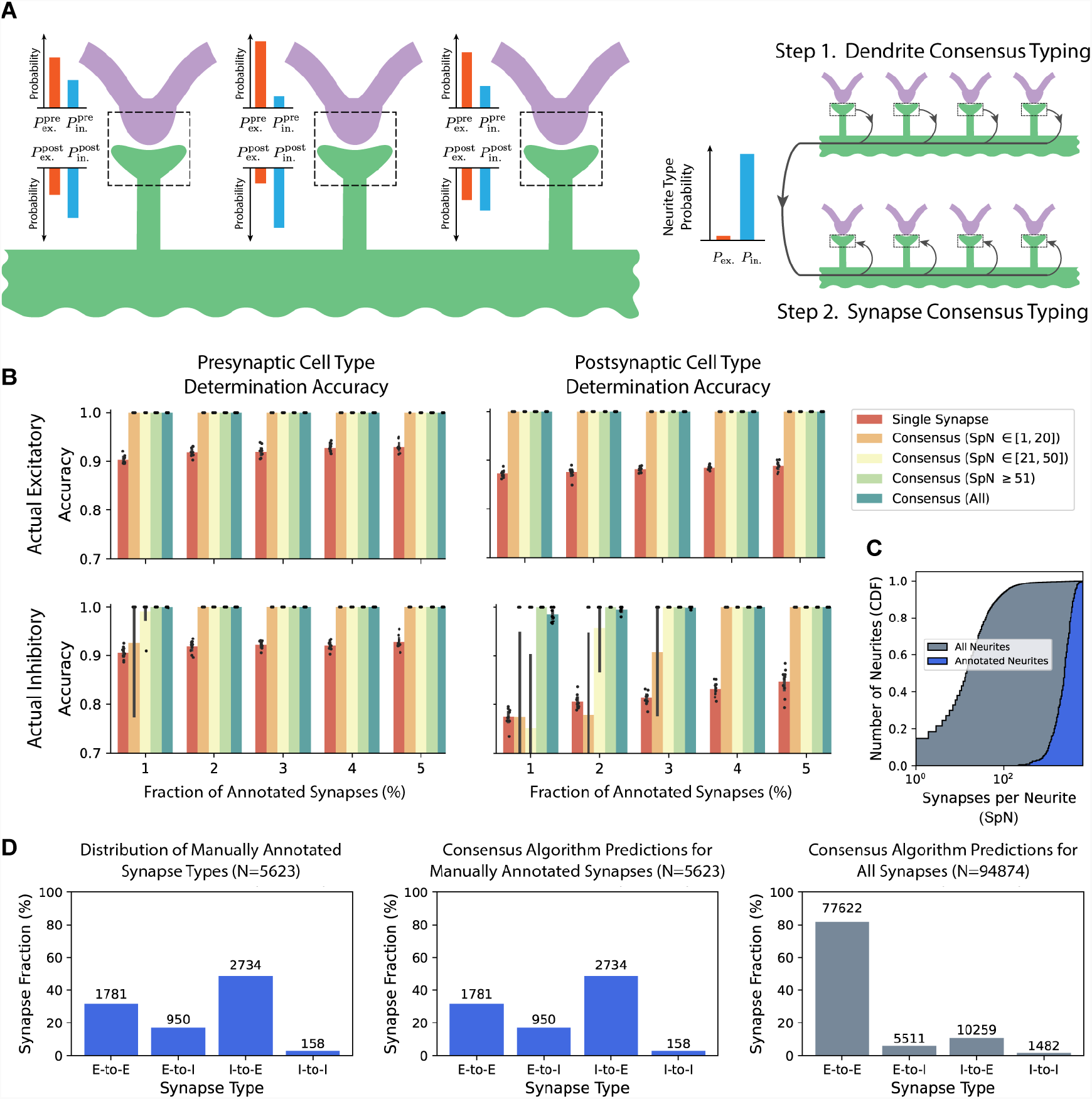
Accurate consensus-based pre- and postsynaptic cell type determination at single-synapse resolution. (A) Left: a schematic illustration of how pre- and postsynaptic cell type probabilities, inferred from single-synapse EM data, can exist for several synapses that involve a single neurite (here, formed onto the same dendrite). Right: the two-step consensus cell-typing algorithm: first, we determine the cell type of each neurite with high accuracy by treating each synapse as a weak classification and ensembling them; next, we propagate the neurite cell type back to the participating synapses (Methods). (B) Consensus-based synapse cell typing cross-validation accuracy, stratified by neurite polarity (pre- or postsynaptic) and actual neuron type (shown in as four separate panels); the number of synapses per neurite (SpN) (shown as differently colored bars); and for different fractions of manually annotated synapses in the dataset, i.e. training data size (x-axis values). The error bars indicate the interquartile range of each accuracy, as obtained by 10-fold train/test partitioning of the annotated synapses. Individual cross-validation runs are shown as dots. Note the dramatic increase in accuracy of consensus calls overall (orange, yellow, and green bars) compared to single-synapse cell type calls (red bars). (C) The cumulative distributions of SpN for all neurites (gray) and for the neurites associated with annotated synapses (blue) in our dataset. The latter group of neurites are distinguished by the fact that their somas are inside the EM imaging volume; as such they are generally more completely reconstructed compared with other neurites and have significantly more synapses. (D) The distribution of pre- and postsynaptic cell types for (left) manually annotated synapses, (middle) consensus cell type calls restricted to manually annotated synapses, and (right) consensus cell type calls over all 94874 randomly chosen synapses from our dataset (L2/3 mouse primary visual cortex). Note the bias in the manual annotations (they are significantly enriched in inhibitory synapses), the perfect agreement between manual annotations and our consensus cell type calls, and the unbiased cell type calls when synapses are chosen randomly from L2/3 primary visual cortex, in good agreement with previous reports of the excitatory to inhibitory synapse ratio.

We validated the accuracy of our consensus cell typing algorithm by randomly censoring the set of annotated synapses we used for training regression models and subsequent consensus calling and then by comparing the inferred consensus pre- and postsynaptic cell types for the held out annotated synapses with their actual values (Methods). We stratified the validation accuracies of consensus calls by actual pre- and postsynaptic cell types, and by several bins of the number of synapses per neurite (“SpN”) to study how many synapses might be required for significant consensus-based improvement in cell typing accuracy compared to cell typing on the basis of individual synapses (Fig. 4B). Furthermore, we repeated this process for multiple levels of censorship to see how much manual annotation is required to train accurate predictive models for the rest of the synapses in the dataset.

We found that excitatory neurites could be typed with virtually 100% accuracy using consensus calling, even at 1% annotation completeness (the lowest level we tested), and with SpN ≤ 20 (Fig. 4B, top panels). We found consensus cell type accuracy to be similar for inhibitory presynaptic neurites (Fig. 4B, bottom left panel), but the accuracies were somewhat lower for inhibitory postsynaptic neurites. We attributed these lower accuracies to the lower prevalence of inhibitory neurons with somas in this EM dataset (14, compared with 364 excitatory neurons with somas). This lower prevalence translates to a relative scarcity of annotated synapses formed onto inhibitory neurons (1108 out of 5623, compared with 4515 synapses onto excitatory neurons; see Fig. 4D, left) and thus, a relatively smaller slice of the training dataset that represents this postsynaptic cell type. Nevertheless, virtually 100% consensus cell typing accuracy was achieved for ≥ 4% annotation completeness.

By studying the distribution of SpN in the full EM dataset (Fig. 4C), we estimated a lower bound for the mean consensus cell typing accuracy over the full ∼3.2 million detected synapses. To this end, we took our least accurate category of single-synapse prediction (postsynaptic inhibitory, p ≃ 0.846 at 5% annotation completeness; see Fig. 4B, bottom right), and assumed all single-synapse predictions to exhibit a similar accuracy. The probability of successful consensus cell typing per neurite for a given SpN can be calculated analytically from the binomial distribution. Explicitly, we take SpN as the number of trials, successful classification probability p = 0.846, and calculate the total probability of having at least (SpN/2) + 1 correct trials. We calculate the mean value of the resulting probability over the empirical distribution of SpN in our dataset (Fig. 4C), and find a mean lower-bound consensus call accuracy of 0.998.

Finally, as a consistency check, we trained the regression models on all of the available annotated synapses and ran the consensus calling algorithm on those same synapses. The resulting consensus cell types were reassuringly in perfect agreement with the actual annotations (see Fig. 4D, middle). Finally, we ran the consensus calling algorithm over the entire set of synapses in order to learn what kind of neurites they were connecting and by extension, to learn about the distribution of synapse types in the dataset. We found that E-to-E synapses dominated the dataset, forming 82% of our sampled synapses (77622/94874). Overall, excitatory synapses (E-to-E plus E-to-I) formed 88% (83133/94874) of synapses in the volume, consistent with previous estimates of the balance of excitatory vs. inhibitory synapses in cerebral cortex^5^; Fig. 4D, right).

### SynapseCLR representations reveal the inherent dimensionality of synaptic structural variation and neuronal subtypes

The accuracy with which our representation regression models can predict complex and fairly abstracted synapse annotations, such as neuronal type, hints at the biologically informed nature of these representations, and encourages us to study the manifold structure of these representations per se. In particular, we are interested to learn about the inherent dimensionality and independent axes of synaptic structural variation, to the degree and granularity that is allowed by the EM data. These data-driven axes of variation will provide a clearer picture of the “isolated synapse”, without dependence on human-derived annotations (many of which strongly co-vary and are not independent, while others might involve information that is not contained in the isolated synapse).

As noted earlier, our UMAP embeddings form two smooth manifolds, one roughly corresponding to E-to-E synapses, and the other corresponding to I-to-E/E-to-I/I-to-I synapses. We used a two-step strategy to learn and interpret the structure of each of these smooth manifolds: (1) We learned directions of variation for each connected component of the UMAP manifold by performing principal component analysis (PCA) on the UMAP coordinates. This procedure provides us with a data-driven set of axes (which we call UMAP-PCs) that describe synaptic structural variation. (2) We study the correlations between the loadings of UMAP-PCs and the annotations in order to interpret these axes of variation in terms of known anatomical measurements. Even though UMAP manifold learning is typically performed in two dimensions, and we have used 2D UMAP as well for visualization purposes, the UMAP algorithm is in fact general and allows learning *n*-dimensional manifold parameterizations. This allows us to repeat the UMAP-PCA analysis for different values of *n* in order to learn higher-order UMAP-PCs. This is a valuable exercise, because it allows us to test whether allowing variation in higher dimensions is necessary for explaining the structure of the representation space. We quantified the relevance of each UMAP-PC in these higher-dimensional spaces according to the amount of variance of UMAP parameterization it explains.

The result of the 2D UMAP-PCA analysis is shown in Fig. 5 (top row: excitatory manifold, bottom row: inhibitory manifold). To guide the eye, the manifolds have been color-coded such that the UMAP-PC 1 axis has a white-to-red gradient and the UMAP-PC 2 axis has a white-to-blue gradient (Figs. 5A,E). Examples from a uniformly sampled 2D UMAP-PC grid (Figs. 5B, F) suggest smooth changes of multiple anatomical features along each direction (see also Fig. S12). The Spearman correlation between the UMAP-PC loadings and the annotations are shown in Figs. 5C and G, together with the ratio of explained variance (EV) by each axis. As Fig. 5C shows, UMAP-PC 1 explained more than 96% of the variance of E-to-E synapses, suggesting that the structural variation of these synapses, by and large, can be explained by a single variable. Moving along this axis is positively correlated with increasing pre- and postsynaptic partner volume; synaptic cleft size (which is limited, but not determined, by pre- and postsynaptic partner volume); and presynaptic mitochondrion presence and volume. The remaining 4% of variation is largely explained by UMAP-PC 2, and extending the analysis to higher dimensions does not change this finding (see Fig. S11A). This second PC correlates with increased presynaptic partner volume; presynaptic mitochondrion presence and volume; and notably, the probability that the presynaptic partner is inhibitory. Indeed, Fig. 3B shows that UMAP-PC 2 is directed toward the region of this manifold where I-to-E synapses are concentrated.

**Figure 5.**
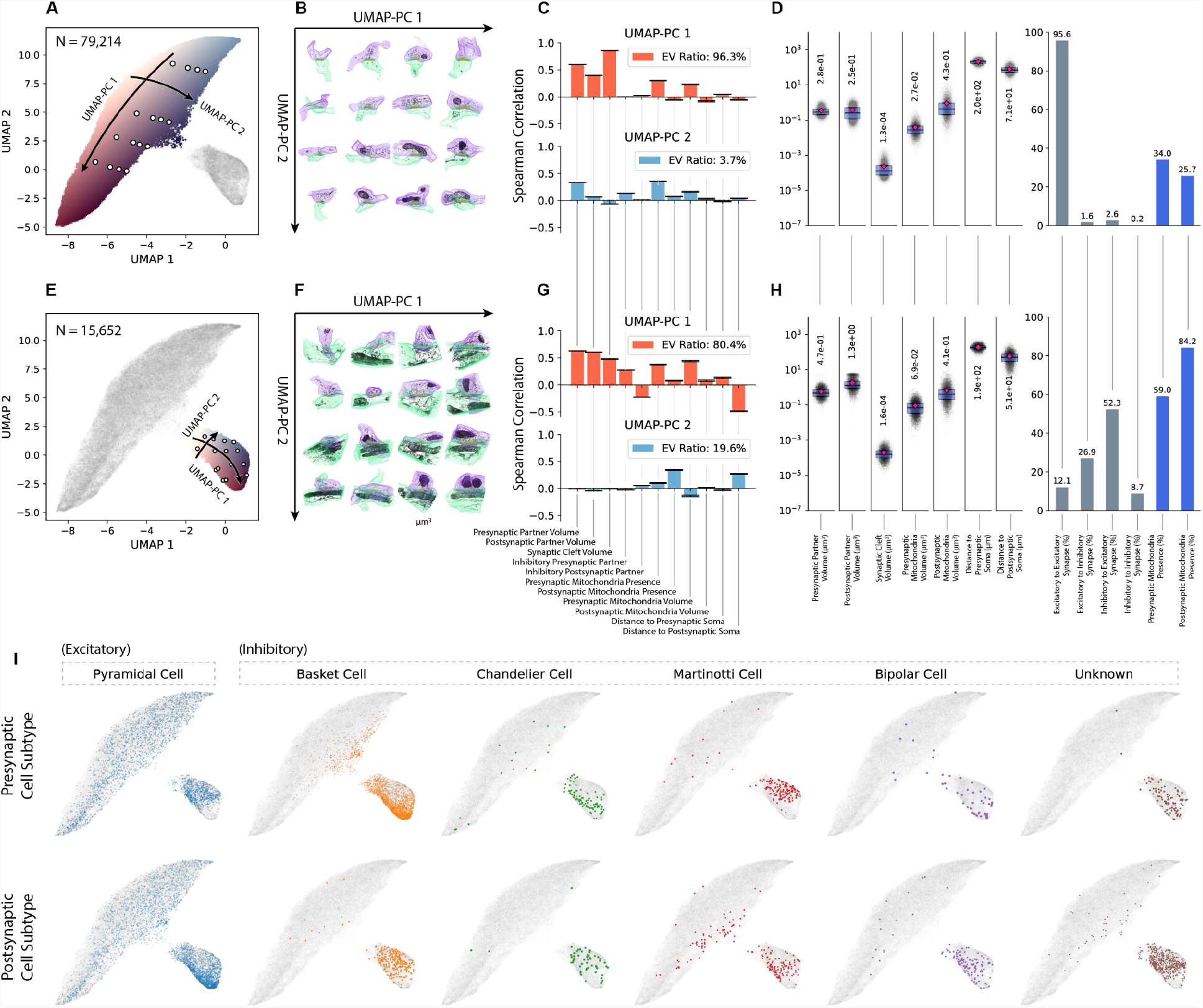
Inhibitory and excitatory synapses form disjoint, low-dimensional, smooth manifolds in the SynapseCLR representation space. (A) UMAP embedding highlighting the excitatory (upper left) sub-manifold, colored by the coordinates of our 2D UMAP-PCA analysis (Methods). The directions of UMAP-PC 1 and UMAP-PC 2 are marked by colored gradients of white-to-red and white-to-blue, respectively. The white circles correspond to the synapses shown in (B). The black arrows are guides for the eye. (B) Synapses along the 2D UMAP-PCA grid of the excitatory sub-manifold, showing visible feature changes including increasing synaptic cleft size along UMAP-PC 1 and presynaptic mitochondria presence along UMAP-PC 2. (C) Spearman correlation of various annotations vs. UMAP-PCs in the excitatory sub-manifold. The legend shows the ratio of explained variance (EV) of the UMAP manifold parameterization by each UMAP-PC. (D) Summary statistics of continuous annotations (left) and discrete annotations (right) for all synapses belonging to the excitatory sub-manifold. Boxes correspond to the inter-quartile range; red markers and numerical values denote the mean. (E) Similar to panel (A) but for the inhibitory (lower right) sub-manifold. (F) Similar to panel (B) but for the inhibitory sub-manifold; note increasing synaptic cleft size and postsynaptic partner volume along UMAP-PC 1, and increasing postsynaptic mitochondria presence along UMAP-PC 2. (G) Similar to panel (C) but for the inhibitory sub-manifold. (H) Similar to panel (D) but for the inhibitory sub-manifold. (I) Pre- and postsynaptic neuronal subtypes shown over a subset of annotated synapses, showing that synapses formed by different neuronal subtypes appear in different parts of the representation space.

In the lower manifold, which contains mostly synapses involving inhibitory neurons, we find that again a large fraction of the variation (80.4%) in representations is explained by UMAP-PC 1. Compared with the excitatory manifold, this reduced amount of variance explained by a single axis indicates that this population of synapses is structurally more diverse. As with the excitatory manifold, repeating this analysis in higher dimensions does not reduce the relevance of the top two UMAP-PCs, suggesting that the inhibitory synapses are largely described by two variables (Fig. S11). In UMAP-PC 1, we observed two relationships of note: first, the probability of the presynaptic partner being inhibitory increases along UMAP-PC 1, whereas the probability of the postsynaptic partner being inhibitory decreases along this axis. Second, the distance to the postsynaptic soma decreases along UMAP-PC 1. The switching of synapse type from E-to-I to I-to-E along this axis, combined with a systematic movement of synapses formed closer to the postsynaptic soma suggested that maybe this axis captured partitioning of synapses formed between different cell subtypes. UMAP-PC 2 correlated with several annotations, but the mixture did not afford us a clear interpretation on the basis of the anatomical properties that might be varying along this axis. However, the variation along UMAP-PC 1 led us to wonder whether UMAP-PC 2 also might be capturing cell-subtype-specific variation in synapse structure. To test this, we highlighted cell subtypes for the subset of annotated synapses that had them (Fig. 5I). All excitatory neurons were labeled “pyramidal” (because no structural subtypes could be easily identified); inhibitory neurons with substantial axons (20) were classified previously as basket, chandelier, Martinotti, or bipolar neurons, or were otherwise labeled as having an unknown subtype^2^. We found that the lower manifold stratified discernibly according to these subtypes. Along UMAP-PC 1, synapses went from being formed by pyramidals onto a mix of inhibitory subtypes (largely basket neurons) to being basket-to-pyramidal synapses (Fig. 5I, top row). Notably, basket neurons form synapses directly onto pyramidal neuron somas, which is consistent with our finding of decreasing distances to the postsynaptic soma in this region (Fig. 3B). Along UMAP-PC 2, inhibitory neurons that formed synapses onto other cell types were remarkably separated according to cell subtype, stratifying into synapses formed by basket, then chandelier, then Martinotti presynaptic neurons. Synapses with bipolar presynaptic neurons also appeared to occupy multiple but specific regions of this manifold; this observation may reflect the fact that there are different structural subtypes of bipolar neuron in this part of the visual cortex^63^. By contrast, the representations of synapses formed onto inhibitory neurons of different subtypes overlap substantially (Fig. 5I, bottom row), indicating that inhibitory dendrites may not differ as much for different inhibitory subtypes at the spatial scale of the synapse.

## DISCUSSION

### Summary

In this work, we have presented a self-supervised contrastive representation learning method called SynapseCLR, an adaption of the SimCLR framework to 3D EM data, and used the method to learn feature representations of synapses in a 3D EM dataset from mouse visual cortex. We established that our representations separated synapses according to both their overall physical appearance and structural annotations of known functional importance. We demonstrated the utility of our methodology for several valuable downstream tasks for the growing field of 3D EM connectomics. These include one-shot identification of defective synapse segmentations, dataset-wide similarity-based querying, and accurate imputation of annotations for unlabeled synapses, using only manual annotation of 0.2% of synapses in the dataset. In particular, we show that neuronal cell types (excitatory vs. inhibitory) can be assigned to individual synapses and highly truncated neurites with accuracy exceeding 99.8%, making this population accessible to connectomics analysis. Finally, we presented a data-driven and unsupervised classification of synaptic structural variation, revealing its intrinsic dimensionality and showing that synapse structure is also strongly correlated with inhibitory neuronal subtypes.

### SynapseCLR representations solve multiple outstanding issues with large-scale 3D EM connectomics data analysis

Our proposed approach to studying synapses is useful for several reasons. First, it is a more scalable, less biased way of making sense of neural circuitry components compared with manual annotation and the conventional supervised machine learning approaches. To learn representations with SynapseCLR, only cell-level segmentations for synapses of interest are needed, as is an understanding of technical sources of variation among images (e.g. variation in pixel intensity, rotation, sectioning artifacts, etc.). This approach reduces the dimensionality of raw synapse images (which can be millions of voxels worth of information) to a few-hundred-dimensional feature space that is, by design, optimally discriminating.

Working with low-dimensional feature representations has two key practical advantages. One is that having a relatively small number of annotated synapses (several thousands) is sufficient to train accurate predictive models without overfitting. We recall that the dimensionality of backbone features (512) is an order of magnitude smaller than the number of annotated synapses (5623), allowing training regression models without overfitting. By contrast, the dimensionality of raw EM image chunks (96 × 96 × 96 ≈ 10^6^) is nearly two orders of magnitude larger than the number of annotated synapses, posing a fundamental challenge for supervised learning. Liberation from intensive manual data labeling has been extensively studied in the context of classifying natural images and is the main motivation behind the development of various self-supervised pre-training regimens^47^. The other advantage of working with low-dimensional feature representations is the possibility of using unsupervised machine learning methods to explore and interpret these representations per se. For example, unsupervised manifold learning using UMAP showed us that representations may split according to neuronal subtype, suggesting that synapses have inherently different structures when formed between different neuron subtypes. Future directions include highlighting additional organelles of interest in explaining synaptic structural variation and linking this to neuronal microcircuits. More broadly, application of unsupervised methods and de-emphasis of human-designed features as afforded by SynapseCLR will be essential for circuit property discovery as EM image volumes continue to grow in size^3,22,42,64^.

Second, our representations highlight not only biologically important variations in synaptic structure, but also non-specific, technically important variations in structure that we did not factor into our augmentations, i.e. segmentation artifacts. Defective synapses can be identified and removed in one shot using only a handful of examples, as shown in Figs. 2 A-F. This characteristic of SynapseCLR representations is reminiscent of few-shot learning approaches in computer vision^53,65^, and is of great value for quality controlling and analyzing large connectomics datasets, where otherwise full, manual inspection of synapses would be required to ensure that the synapses being studied are not contaminated.

Third, SynapseCLR representations capture features that in particular define pre- and postsynaptic cell type (E vs. I; Fig. 2I) with high accuracy. By ensembling single-synapse predictions over individual neurites, we were able to to assign highly accurate cell types for the many highly truncated neurites in this EM dataset that formed most of our unannotated synapses (Fig. 4). This capability provides huge value to the field of connectomics, because usually cell types can only be assigned to mostly complete neurons (or glia) in a dataset which comprise a small minority of neurites (here, 400 of 8 million, or 5×10^−3^%, of objects) and the majority of objects are left out of the analysis. By running imputations and making consensus calls on neurites that form 2 or more synapses, we estimate that we are able to put ∼378000 of them back into our analysis with ∼100%-accurate cell types (roughly 30% of the 1.27 million neurites that form putative synapses in the volume, and 4.7% of the 8 million neurites in the volume), significantly enhancing future studies of cell-type-specific connectivity patterns.

### Appropriate augmentation design is critical for finding meaningful self-supervised representations

The learning paradigm used in SynapseCLR is designed to retain the most robust discriminating visual features that remain invariant under a specified set of augmentations, while distilling out non-invariant visual data. Thus, in order for SynapseCLR to generate biologically meaningful synapse representations, augmentation choice is critical: augmentations should be designed to capture as much technically-sourced image variation as possible, and to avoid capturing biological variations. We motivate our augmentation choices in great detail in the Methods section. As a concrete example of the way insufficient augmentation can degrade representations, we trained SynapseCLR a second time with a modified type of rotation. With our original approach, where full SO(3) rotations were allowed (giving the results in Figs. 2-5), we see that the representation space is indeed invariant to synapse orientation angle (Fig. S7A, C) while successfully capturing variation in other structural properties (see Results). (One exception is the subset of E-to-E synapses that occupy the upper right region of the upper manifold; this may be indicative that synapse orientation is highly conserved for synapses formed by some excitatory cell subtypes; see below). In contrast, when we restricted augmentations to discrete octahedral rotations only, the resulting representations were dominated by synapse orientation (Fig. S7B). As a control demonstrating that our augmentation procedure is functioning properly, the number of blanked sections was varied in augmentations for both runs, and in both the resulting representations had no dependence on the number of missing sections.

### SynapseCLR representations suggest new directions for future studies of neuronal circuit connectivity

The representations generated by SynapseCLR show that structural variations among synapses correspond to notable annotation properties that should be further investigated. First, by studying the axis of greatest variation for E-to-E synapses (the long axis of the upper manifold; Fig. 5C, UMAP-PC1) we see that the primary way in which synapses differ along this axis is by cleft size, and it appears that synapses with larger clefts have more presynaptic mitochondrial presence and less postsynaptic mitochondrial presence than smaller synapses. These relationships could be a marker for E-to-E synapses that are being potentiated: when a pre- and postsynaptic cell repeatedly fire in a fixed temporal pattern, with presynaptic firing followed tens to hundreds of ms later by postsynaptic firing (this is also called “synchronous” firing), synapses are thought to undergo changes that mediate more efficient signal transmission^66,67^. One such change for excitatory neurons over longer times (roughly hours or longer) is addition of AMPA receptors^67^; this would correspond to a larger synaptic cleft in an EM image, since cleft size is typically measured as the volume of the postsynaptic density (the net of architectural proteins and receptors positioned on the postsynaptic side of the synapse^31^). Mitochondria may also have a role in promoting synaptic potentiation: in addition to providing cells with energy (via oxidative phosphorylation that produces ATP), mitochondria buffer Ca^2+^ out of surrounding cytosol^46^. Ca^2+^ is essential for the release of vesicles containing neurotransmitters from the presynaptic cell, and one consequence of reduced levels of presynaptic Ca^2+^ is reduced spontaneous vesicle release. Mitochondria-mediated removal of presynaptic Ca^2+^ suppresses such “asynchronous” activity and promotes synchronous firing leading to potentiation^68^. In the postsynaptic region, mitochondria are the primary energy source responsible for local translation of mRNA into proteins^46^. It has been proposed that enhanced postsynaptic mitochondrial presence may be a signature of synaptic downscaling, which is in turn a consequence of synaptic potentiation: broad postsynaptic downscaling allows potentiated synapses to take larger weights in signal transmission without triggering runaway activity (epileptic events)^69,70^. Future work could focus on determining whether these distinctions do in fact indicate a subpopulation of potentiated E-to-E synapses.

Second, as highlighted by our UMAP-PC analysis, representations appear to separate synapses not only by pre- and postsynaptic cell type, but also by cell subtype (as we see from our synapses involving inhibitory neurons; Fig. 5I). This finding is important because it indicates that neurites can be assigned cell subtypes based on the structure of the synapses they form. In the future, annotations of these synapses could be used to impute cell subtypes for neurites in EM datasets and thus add a layer of specificity to connectivity analyses. This type of information could further build out the picture of cortical connectivity, by highlighting the frequency of various cell connection types and describing the synapse properties of those connections.

The finding that inhibitory neurons can be subtyped based on their synapses begs the question of whether the same can be said about excitatory neurons. As shown in Fig. S7, the E-to-E manifold does possess a region where synapse orientation is important (the upper-right region). This region could either correspond to synapses involving a specific compartment of an excitatory neuron (pyramidal cells have highly stereotyped dendritic morphologies, including apical dendrites that point radially and basal dendrites directed 45 degrees away from the apical axis), or it could correspond to a specific subtype of excitatory neuron. Further work to identify annotations that might identify excitatory cell subtypes (like compartment labels or cortical layer labels, e.g. L2/3 vs. L5/6) would be useful for learning more about excitatory subtypes.

Additionally, the finding that E-to-E synapses may separate based on potentiation state begs the question of whether this is true for inhibitory neurons. Indeed, this and many other facets of the synapses in this dataset can be investigated by identifying additional structural correlates of function. For example, synaptic vesicles (both docked and free^71^) and endoplasmic reticulum^39^ are two organelles already known to be essential for synaptic function that could be annotated to help interpret SynapseCLR representations. One of the salient characteristics of contrastive learning is the implicit inversion of the data generating process (here, meaning aspects of cellular function) by learning disentangled representations at the projection head^72,73^. Here, a disentangled feature means one that controls a specific aspect of the synapse structure while remaining inside the data manifold and leaving other aspects intact (i.e. being an independent knob). We have empirically demonstrated this emergent property for SynapseCLR representations in Fig. S8A, where projection head features are found to be largely independent of one another. One may speculate that disentangled visual features of synapses may correspond to biological function. Indeed, we find that projection head PCs 1 and 8 control cleft size and mitochondrial presence (though not independently), both features being of known functional significance. Projection head PC 3, on the other hand, while being largely uncorrelated with our available manual annotations, seems to be responding to synaptic vesicles upon visual inspection. More generally, we speculate that relating SynapseCLR representations to structural correlates of function, paired with visual assessment of synapses showing high and low activation of these features, may reveal novel anatomical properties of synapses tied to their function.

SynapseCLR representations themselves can be enhanced by several future endeavors. First, training on larger input image chunks could allow for more relevant information over larger spatial scales to be encoded. Such information could include how synapses may vary with distance from cell somas (in this work we trained on images spanning a few μm, whereas dendrites and axons extend for 100s of μm and up to 400 μm in this dataset^4^). It could also allow for more informative encoding of mitochondrial features by capturing more of the mitochondria near synapses: in our image chunks, mitochondria were often truncated because they were either much larger than the volume (true for most postsynaptic mitochondria), or because they were slightly farther from the synaptic cleft than the image chunk boundaries (a source of noise for both pre- and postsynaptic mitochondria). Similarly, training with a larger network to produce SynapseCLR representations could lead to more detailed representations, as has been shown by past work^47,49^.

SynapseCLR can be readily used for learning representations from other objects in an EM dataset besides synapses. Correlating synapse representations with the representation of the surrounding neuropil is an attractive future direction for revealing potential modulation of synapse structure by nearby processes (e.g. microglia and other glial cell types are known to modulate synaptic function^37,56^).

Looking farther into the future, generation and interpretation of unsupervised representations in EM could be further enhanced by adding multimodal information, such as focused, higher-resolution EM; molecular image labels through immunohistochemistry or by other means; or activity-based images using techniques such as Ca^2+^ imaging. Specific targets for either input labels or annotations could include electrical synapses^74^ or more subtle synaptic organelles like multivesicular-endoplasmic reticulum bodies^75^. Given that transcriptional states are strongly correlated with morphological features in cells and tissues^76–79^, we believe that informative morphological representations such as those produced with SynapseCLR may help form a more complete picture about the connection between structural connectivity and the underlying molecular mechanisms.

## ACKNOWLEDGEMENTS

The authors thank Stephen Fleming, Luca D’Alessio, Nicholas Barkas, Anthony Philippakis, and William Silversmith for insightful discussions at various stages of this project. A.M.W. acknowledges financial support from NIH NIMH BRAIN Initiative award number RF1MH123400; IARPA contract number D16PC0005; NIH NIMH award number T32MH065214. M.B. acknowledges financial support from Data Sciences Platform (DSP), Broad Institute.

## METHODS

### EM Source Dataset

Our 3D EM image volume was a reconstruction from layer 2/3 (L2/3) of primary visual cortex in a young-adult (postnatal day 36), male mouse (from the IARPA MICrONS project; see https://www.microns-explorer.org/phase1). This volume was built from ∼2250 EM images, each 40 nm thick and with resolution 3.58 × 3.58 nm^2^, and measured 240 × 140 × 90 μm^3^ in total. We made use of 3 segmentation layers that were also released with this dataset: one layer segmenting the EM images into its constituent cells; one identifying the active regions of chemical synapses (called “synaptic clefts”) in the volume; and one showing all the mitochondria present in the volume. We also used skeletons for 397 excitatory and inhibitory neurons with their somas in the volume. These skeletons had been proofread and smoothed to allow for calculations involving distances along the dendrites or axons of these neurons.

### EM and Segmentation Mask Image Chunk Generation

To generate image tensors for training, we pulled the list of synaptic clefts published with the MICrONS L2/3 mouse primary visual cortex dataset^80^. This list contains 3.2 million rows of putative synapses and a subset of 6761 synapses that had been previously proofread by multiple experts. These synapses were formed between excitatory and inhibitory neurons that had their somas contained inside the dataset and were used for multiple analyses^4,35,81^.

We then sampled additional synapses to generate a targeted training dataset of 100000 synapses, by randomly sampling from the rest of the synapse list (which contained synapses scattered effectively uniformly throughout the EM dataset). For every synapse we chose, we downloaded a 256 × 256 × 256-voxel EM image chunk at MIP 1 (7.16 × 7.16 × 40 nm^3^/vx) from the public data repository, and only added it to our input image if <10% of the image chunk was missing (i.e. masked out to zeros in order to prevent section collection or imaging artifacts from affecting image volume post-processing). If a chunk had more than 10% of its volume masked out, it was discarded from the input set and a different synapse was selected and checked for suitability.

If the synapse met our criteria for EM masking, we also downloaded chunks of the same size and centered at the same location from the segmentation, synaptic cleft, and mitochondrial layers, to be used for input tensor generation and, for synapses connecting neurons with somas in the volume, annotation population. Input image data was downloaded for all synapses using CloudVolume.

For the segmentation layer, we restricted our download to only the pre- and postsynaptic neurons forming the current synapse, using the synapse table provided with this dataset. If this process failed for any of the synapses for which annotations were possible, we discarded it from our training set. Because we also used mitochondrial segmentations for our annotation-based analysis, we also discarded synapses where the mitochondrial segmentation was not available. This process resulted in a training dataset consisting of 5664 annotated and 91994 unannotated synapses.

Keeping in mind that we ultimately wanted to work with isotropic images, we kept only the 52 central z-sections that covered roughly the same physical linear dimension as x and y (i.e. 52 sections at 40 nm/px = 2080 nm, 256 pixels at 7.16 nm/px = 1832 nm). During data augmentations, we then upsampled our z pixels to roughly our match x-,y-resolution (see below).

### Volumetric Data Augmentation

We constructed a set of intensity-, masking-, and affine-transform-based augmentations to use for training our network on synapse image data (Fig 1C). Each transformation is defined, first, by a probability parameter, which fixes the probability at which it will be included in an instance of an augmentation, and second, by hyperparameters that define the exact orientation, shape, or magnitude of the transform.

We implement augmentation transformations in PyTorch to leverage GPU acceleration. Transforming volumetric data is considerably more time consuming than 2D images and the conventional CPU-based approach to data augmentation results in considerable slowdown and wasted GPU time. The overall data flow for augmenting a batch of *n* volumetric image chunks is as follows:

1. The volumetric intensity data is transformed to [0, 1] value range and is represented as a batch of 3D tensor of size 256 × 256 × 52, and the binary segmentation masks (presynaptic process, synaptic cleft, postsynaptic process) are represented as a batch of 3-channel 3D binary tensor of size 256 × 256 × 52. We note that at this stage, the representation is anisotropic: each voxel corresponds to a physical volume of 7.16 × 7.16 × 40 nm^3^.
2. A first series of intensity-based and mask-based transformations are randomly sampled and applied to the intensity and the mask tensors, respectively. These transformations are described below.
3. An affine transformation composed of a random 3D rotation, random center displacement, central cropping, and a fixed anisotropic scaling is applied to both intensity and mask tensors. Central cropping and anisotropic scaling transformation plays three roles: (1) reverting the unequal sectional and in-plane tissue imaging resolution and ensuring that the final voxels are isotropic at 7.16 nm/px a side; (2) shrinking the field of view from ∼2 μm^3^ (256 px) to ∼1.5 μm^3^ (192 px) in order to reduce the external context; (3) downsampling the resolution to the final desired output resolution of 96 × 96 × 96 voxels, with each voxel corresponding to a physical volume of 14.3 × 14.3 × 14.3 nm^3^. We implement this series of transformations efficiently by leveraging PyTorch’s Spatial Transformer Network (SPN) capabilities as a single operation^82^.
4. A second series of intensity-based and mask-based transformations are randomly sampled and applied to the isotropic intensity and the mask tensors, respectively. These transformations are described below.
5. Finally, the 3-channel mask tensor is merged to a single binary mask via a voxel- and bit-wise OR operation. Peripheral voxels that may be disconnected from the synaptic cleft are identified and removed using cc3d package^83^ by traversing the 26 nearest neighbors of each voxel. The intensity channel is multiplied with the resulting “active zone” binary mask, z-scored using dataset-wide mean and standard deviation, and returned.

We will describe each transformation and our choice of hyperparameters below.

#### Intensity-based Transformations

##### Sectional cutout

A common technical artifact of 3D EM datasets generated via serial sectioning is the random missingness of parts of many scanned sections due to tissue sectioning or scanning issues (e.g. thin tissue deformation, warping, folding). A typically employed quality control measure is to identify and blank out affected regions along the fold line. Since the blanked out regions can be tens of microns in width, these partially missing regions appear as completely missing sections when sampling micron-sized 3D volumes (e.g. a synapse) from the reconstructed dataset. Such a prominent technical artifact may be easily exploited by the representation learner to trivially solve the contrastive objective without learning a deeper structural feature representation. We estimated a typical synapse to have 1-2 missing sections in our dataset. To disincentivize the model from incorporating such artifacts into the representation space, we randomly select *n*_*cutout*_∼ *Poissson*(10) sections and blank them out. This transformation is applied before the affine transformation and with probability 0.5.

##### Sectional intensity distortion

The EM dataset is normalized appropriately to minimize sectional intensity variations. The normalization strategy implemented by the original authors, however, is not locally adaptive and thus, we estimated approximately 10% sectional intensity variation. To desensitize the representations to such such technical variations, we apply following transform stochastically to every local z-section:

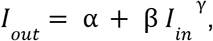

where *I*_*in*_ and *I*_*out*_ denote the input and output z-sections, respectively. The transformation is applied pixel-wise, and for each z-section, α, β, and γ are independent and identically distributed (i.i.d.) random variables sampled as follows:

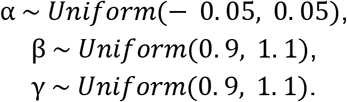

The output intensities *I*_*out*_ are clipped to range [0, 1] to prevent under- and overflows. This transformation is applied before the affine transformation and with probability 1 (always).

##### Global intensity distortion

A similar reasoning applies to technical intensity variations at the volumetric level, i.e. presence of intensity variations at the scale of tens of microns due to varying tissue quality, scanning batch, etc. As a remedy, we apply the same transformation as above to the entire 3D volume:

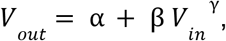

where *V*_*in*_ and *V*_*out*_ denote the input and output volumes, respectively. The transformation is applied pixel-wise, α, β, and γ are sampled like before. In contrast to sectional distortion, α, β, and γ are not independent across z-sections and are sampled once for the entire volume. The output intensities *V*_*out*_ are clipped to range [0, 1] to prevent under- and overflows. This transformation is applied before the affine transformation and with probability 1 (always).

##### Pixel noise

Independent pixel noise is a ubiquitous source of noise in any imaging. We apply both normal additive and log-normal multiplicative independent noise to every voxel:

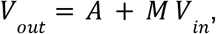

where *A* and *M* (positive) are 3D additive and multiplicative i.i.d. noise tensors with the same dimensionality as *V*_*in*_. We estimated the pixel noise level to be approximately 1% to 5% by comparing the intensity of adjacent pixels. Moreover, we observed variation in the level of noise at the scale of tens of microns (likely due to varying tissue condition). We model this using a simple two-hierarchy noise sampling procedure:

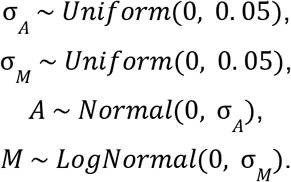

The output intensities *V*_*out*_ are clipped to range [0, 1] to prevent under- and overflows. This transformation is applied before the affine transformation and with probability 1 (always).

##### Gaussian blur

Another source of imaging variation is blurring, which in EM datasets, stems from variation in tissue quality and dye diffusion. While blurring is not considered a significant issue for the scanning EM imaging modality (as opposed to e.g. fluorescence microscopy), we reasoned that its inclusion as an augmentation may help decrease the reliance of the model on high-resolution features. We apply Gaussian blur to each volume with probability 0.5 after the affine transformation. If blurring is to be applied, we randomly select a blurring radius σ_*blur*_ from 3 options with equal chance: 0.25, 0.5, or 0.75 (in the final units of 14.3 nm), generate a normalize 3 × 3 × 3 Gaussian 3D convolution kernel *K*, and apply the 3D convolution to the intensity tensor:

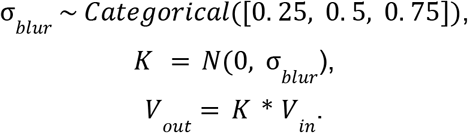

#### Mask-based Transformations

##### Peripheral cutout

We encourage the model to attend to synapse “active zone” by disincentivizing it to use peripheral intensity data using three different transformations:

1. *Corner cubic cutout:* We subdivide the synaptic volume in thirds along each axis, randomly select one of the 26 cubes neighboring the central region, and blank it out with probability 0.5.
2. *Random cubic cutout:* We randomly select and blank out 2 cubic regions, each ranging in dimensions from 12 × 12 × 12 voxels to 36 × 36 × 36 voxels, with probability 0.25. We protect the synaptic cleft region from being selected and cropped.
3. *Center crop:* We select a cubic region around the synaptic cleft with linear dimensions ranging from 50% to 70% of the entire box, with a maximum center displacement of 20%, and we blank out the exterior region with a probability of 0.25.

These transformations are applied after the affine transformation in the final 96 × 96 × 96 voxels space.

##### Segmentation mask inflation

We disincentivize the model from exploiting potential imperfections of the segmentation masks by inflating (dilating) the segmentation masks with randomly chosen inflation radii. We implement the operation by convolving the binary masks with a cubic dilation structure as follows:

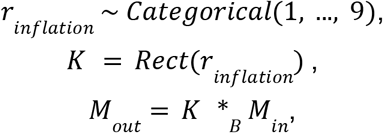

where *r*_*inflation*_ is the inflation radius, *Rest*(*r*_*inflation*_) is the cubic dilation structure, i.e. a 3D kernel with shape (2*r*_*inflation*_ + 1)^3^ with 1 everywhere, *M*_*in*_ and *M*_*out*_ are the input and output binary segmentation masks, and *_*B*_ is the binary convolution operator.

#### Data Augmentation in Batch Mode

When augmenting a batch of images, we implement two modes of operation: (1) the “coupled” mode, where the same randomly composed augmentation transformation is applied to every image in the batch, and (2) the “decoupled” mode, where a different randomly composed transformation is generated and applied to each image. In theory, the “coupled” mode leads to both harder positive and negative examples: for any given image in a batch of *n* images, the same transformation is applied to *n - 1* other images, while also the matching image (second in pair) shares the same augmentation transformation with *n - 1* negative images. As a result, the contrastive loss cannot be trivially minimized by learning the signature of augmentation transformations and exploiting those signatures to rule out potential positives or negatives. In our preliminary experiments, we observed evidence supporting improved representations when training the model using coupled augmentations (e.g. a more structured UMAP embedding) in agreement with our theoretical reasoning. The results shown in this paper are all generated using models trained using coupled augmentations, a choice that deviates from the original SimCLR paper^47^. A systematic empirical study of the impact of different augmentation strategies on the quality of representations is a valuable future research direction.

#### Feature Extraction Mode

A critical part of our raw data preprocessing, namely bilinear upsampling of the axial resolution to match the in-plane resolution, and downsampling to the final 96 × 96 × 96 resolution (which is an allowed input dimension for 3D-ResNet-18 CNN architecture), is done on the fly and as a part of our fast GPU-based data augmentation. Following successful model training, we wish to process each synapse with the trained CNN and extract representations from different layers (backbone, projection middle, projection head). To this end, we simply disable all intensity-based, mask-based, and affine transformations except for the anisotropic scaling component of the affine transformation step. In this mode, the augmentation module operates in a fully deterministic mode and simply acts as a fast data resolution transformer.

### SynapseCLR Loss Function and Training Schedule

We closely follow the SimCLR methodology developed for natural images^47^, with two modifications: (1) We use a 3D-ResNet18 CNN as the backbone instead of 2D-ResNets; (2) We replace the SimCLR data augmentation transformations (relevant to 2D natural images) with the 3D EM-specific transformations discussed in the previous section. In this section, we will provide a brief description of our adapted framework (“SynapseCLR”), including the neural network architectures, loss function, and training schedule.

#### The SynapseCLR Framework: Network Architecture and Loss Function

Each iteration of SynapseCLR training starts with a minibatch of *N* randomly selected images, *x*_*i*_ for *i* = 1, …, *N*. Next, we generate two random views (“augmentations”) of each image by composing a series of randomly selected transformations, *t*_*i*_ for *i* = 1, …, 2*N*, from a predefined set of transformations. These transformations are applied to the images in the minibatch and the results are interleaved in even and odd pairs:

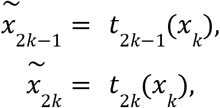

for *k* = 1, …, *N*. The augmented images are first processed by the CNN backbone *f*(.), followed by the projection head *g*(.), in order to generate a final representation for contrasting and matching:

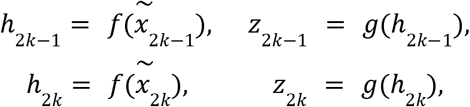

for *k* = 1, …, *N*. The role of the projection head is to further process the CNN features and embed them in a space amenable to contrasting and matching using one’s similarity metric of choice (see below). The architecture of the 3D-ResNet18 and projection head we have used in our implementation is shown in Fig. S1A. Our ResNet backbone is largely a 3D adaptation of the original 2D ResNet^55^ with a minor modification of the network stem: given that our input images are already downsampled to 96 × 96 × 96, with each voxel corresponding to ∼14.3 nm in physical space, we reasoned that a max pooling operation commonly in the network stem would result in loss of high frequency visual information and therefore bypassed it.

The SimCLR contrastive training objective is to encourage two views of the same image to be as similar as possible, i.e. *h*_2*k*−1_ ≈ *h*_2*k*_, while maintaining dissimilarity to both views of every other image in the minibatch. This is achieved by calculating and summing a normalized temperature-scaled cross entropy (“NT-Xent”) loss function for each image in the augmented minibatch:

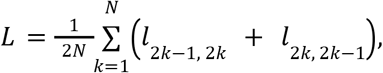

where:

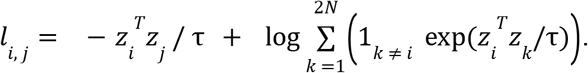

Here, τ is the temperature hyperparameter that sets the typical range of repulsive and attractive forces in the representation space. It has been empirically shown that τ = 0. 5 is a robust choice independent of the learning rate and minibatch size^47^.

Prior to model training, the projection head representations *z*_*i*_ are largely uninformative. As such, the cosine similarity values *z*_*j*_^*T*^*z*_*j*_ appearing *l*_*i,j*_ are drawn from the same distribution for both positive and negative pairs. As a result, *l*_*i,j*_ ≈ log (2*N* − 1) independently of *i* and *j* and the initial value of the loss function is expected to be *L*_*init*_ ≈ log (2*N* − 1). As model training proceeds, we expect the appearance of a separation in cosine similarity between positive and negative pairs, i.e. *z*_*j*_^*T*^*z*_*j*_ − *z*_*j*_^*T*^*z*_*k*_ ≈ Δ > 0, for *i* and *j* being two views of the same image and *k* being a different image. Consequently, we find *l*_2*k*−1, 2*k*_ ≈ *l*_2*k*, 2*k*−1_ ≈ log (2*N* − 1) − Δ / τ. Therefore, a drop in the loss function *L* from its initial value by an amount δ*L* implies the emergence of a separation Δ ≈ τ δ*L* in the cosine similarity between positive and negative pairs.

#### Initialization and Training Schedule

We initialized the convolutional layers using Glorot-scaled Gaussian weights^84^ truncated to [-2, 2]. Weights and biases of the linear layers were initialized to i.i.d. draws from N(0, 0.01). The 3D-ResNet18 and projection head contain ∼ 33.4M and ∼ 329K trainable parameters, respectively. We used the Adam optimizer with an initial learning rate of 2e-4 with a cosine decay to 2e-5 during 100 training epochs. After 100 epochs, we reset the learning rate back to 2e-4 and perform another 100 epochs of contrastive training. We used 4x NVIDIA A100 GPUs each with 40 GB of HBM2 RAM. With 160 GB of GPU RAM, we could upload, augment, and perform a complete forward and backward pass on a minibatch size of 192 synapses (48 synapses/GPU). Training for 200 epochs took approximately two weeks. The NT-Xent loss function is shown in Fig. S1B and is largely stationary after restart, suggesting convergence.

### Annotations for Verified Synapses

For EM image chunks of verified synapses, we also annotated several properties that previous studies indicate may be related to synaptic function and thus potentially also of structure:

- **Presynaptic cell type:** Assigned as excitatory or inhibitory from the soma valence table provided in https://www.microns-explorer.org/phase1.
- **Postsynaptic cell type:** Also assigned as excitatory or inhibitory from the soma valence table.
- **Synaptic cleft size:** The number of voxels forming the synaptic cleft, as reported in the synapse table from the dataset.
- **Distance from synaptic cleft to the soma of the presynaptic neuron:** Computed by finding 1) the node in the presynaptic neuron skeleton closest to the synaptic cleft centroid, and 2) the node of the presynaptic skeleton corresponding to the soma centroid. We then used the networkx package in Python to convert the presynaptic skeleton to a graph, and used the shortest_path_length algorithm to estimate the distance between these two locations.
- **Distance from synaptic cleft to the soma of the postsynaptic neuron:** Computed using the same process as above with the postsynaptic neuron.
- **Presence of presynaptic mitochondria:** Mitochondrial segmentations within 5 μm of annotated synapse centroids were manually proofread. If mitochondrial segmentations overlapped with the presynaptic segment in an annotated synapse image chunk, this value was set to 1; otherwise it was set to 0.
- **Presence of postsynaptic mitochondria:** Same as above but mitochondria overlapping with the postsynaptic segment were used.
- **Volume of presynaptic mitochondria:** The number of voxels of mitochondrial segments overlapping with the presynaptic segment in the synapse image chunk.
- **Volume of postsynaptic mitochondria:** Same as above, but for mitochondrial segments overlapping the postsynaptic segment.

### Gaussian Process (GP) Regression for Annotation Imputation

In this section, we describe our methodology for probabilistic imputation of annotation for unlabeled synapses. Conceptually, we wish to use SynapseCLR representations as covariates (“input space”) and predict the annotations (“target space”) using available manually annotated synapses. Previous authors have used linear classifiers and regressors for this task^47^. Here, we propose to use Gaussian process (GP) models instead. GP has two key advantages compared to linear models: (1) linear models can be thought GPs with linear kernels; using appropriate nonlinear kernels may prove to be an effective way to improve the accuracy of predictions; (2) GP models are naturally endowed with a Bayesian posterior probability distribution, allowing us to additionally model of the credibility of our predictions.

#### Annotation Preprocessing and Class Balancing

To facilitate GP training, we log-transformed and z-scored all continuous annotations to compress and standardize their dynamic ranges. This transformation is invertible, allowing us to revert to the original scale after performing imputation. In light of the presence of significant sampling bias in our available annotations (e.g. there are more inhibitory synapses than excitatory synapses in the annotation set; see Fig. 4D), it is crucial to perform class balancing on the GP training dataset. To this end, we randomly resampled an equal number of synapses from each binary class 5 times and trained separate GP models on each rebalanced dataset. We observed reassuringly negligible differences in predictions and reported ensemble average of the GP posterior predictions.

#### Cross Validation and Accuracy Metrics

For an unbiased assessment of the accuracy of the GP fits, we split the annotated dataset into 90% training and 10% testing sets, fit the GP on the training set and reported accuracy figures on the testing set. We repeated this procedure 5 times with different random splits to estimate the confidence level of the reported accuracy figures. We report the prediction accuracy of continuous variables in terms of explained variance (EV) percentage over the test set defined as follows:

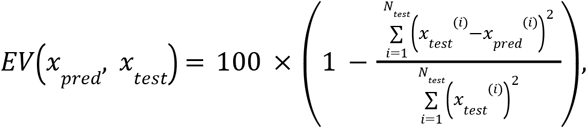

where *x*_*test*_^(*i*)^ and *x*_*pred*_ ^(*i*)^ the denotes expected and predicted value of a specific annotation for the *i*’th synapse in the test set. For binary traits, we report the area under the receiver operating characteristic (precision-recall) curve (AUC), calculated using sklearn.metrics.auc function implemented in scikit-learn (Ref).

#### GP Fitting and Hyperparameter Optimization

We leveraged the GP functionality available in the Pyro probabilistic programming language^85^. In particular, we used the variational sparse Gaussian process (VSGP) flavor using a mean-field posterior for speed and scalability.

We trained separate GP regressors and categorical classifiers for each continuous and discrete annotation. In principle, better fits might be achieved via multi-task training (co-kriging), however, we did not attempt that here. We trained separate GP models on 512-dimensional CNN backbone, 512-dimensional projection middle, and 128-dimensional projection head features as covariates. In each case, we initialized the VSGP inducing points using k-means. We used the Adam optimizer with an initial learning rate of 1e-3 with cosine annealing to 0 during 20,000 training epochs. We considered performing PCA dimensionality reduction on the covariates before fitting, however, we found inferior performance in every case (see below).

We used the previously described cross-validation strategy and accuracy metrics to optimize: (1) the number of VSGP inducing points, and (2) the choice of GP kernel. A summary of the experiments are shown in Table S1, pointing to the following general conclusions:

1. Using the 512-dimensional CNN backbone features as covariates provided the highest prediction accuracy for all of the annotations (see the first group of rows in Table S1)
2. PCA dimensionality reduction of covariates, regardless of number of kept PCs, degraded the accuracy of predictions (see the second group of rows in Table S1).
3. We considered the following kernels:
  a. linear kernel with wihte noise:

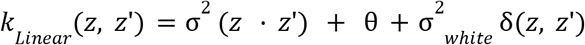
  b. Laplace kernel with white noise:

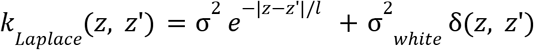
  c. Radial basis function (RBF) kernel with with noise:

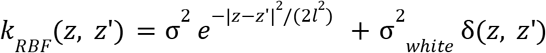

The best fits were obtained using the RBF kernel with white noise and with a fine-tuned number of inducing points. The latter effectively acts as a regularization by controlling the model complexity, allowing us to prevent overfitting which is made possible using the more flexible class of functions reproduced by the RBF kernel (see the third group of rows in Table S1).

Finally, Fig. 3A shows the accuracy of best GP fits obtained by following the same comprehensive hyperparameter optimization strategy using features extracted from a random CNN baseline and MedicalNet, a general-purpose pretrained 3D-ResNet18 CNN. We found that SynapseCLR features lead to significantly higher GP imputation accuracy figures.

### Ensembling and Consensus-Based Cell Typing

We showed that SynapseCLR representation learning followed by fitting GP classifiers can reliably predict pre- and postsynaptic cell types for individual synapses (0.975 and 0.940 AUC, respectively; see Fig. 3A). As mentioned in the main text, we may further exploit the fact that a typical synapse-forming neurite in our dataset contains typically tens of synapses. This figure reaches thousands of synapses for unfragmented neurites. A highly effective way to boost the accuracy of several weak classifiers is by ensembling, i.e. by taking the consensus of all weak classifiers. In the present context, single-synapse cell type predictions can be thought of as weak classifications. By appropriately chaining weak predictions over synapses sharing the same neurite, the neurite-level cell type prediction error decreases rapidly and reaches practically noiseless limits. The consensus cell type of each neurite can then be propagated back to the participating synapses (see Fig. 4A). We carry out this procedure in two steps:

#### Step 1 (Neurite Cell Typing)

We use the 3D segmentation and skeletonization information available to us to compile a list of synapses formed onto each neurite. We proceed to calculate a “consensus” cell type probability for each neurite. To this end, we use a minimal Bayesian model in which the identity of a given neurite determines the pre- and postsynaptic cell type probabilities *p*_*n*_ of individual synapses form onto it:

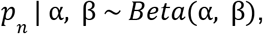

where *n* = 1, …, *N*_*syn*_ enumerate the synapses formed onto the neurite. We assume *p*_*n*_ = 0 and *p*_*n*_ = 1 for excitatory and inhibitory cell types, respectively. We assume *p*_*n*_ are i.i.d. conditioned on α and β, the concentration parameters of a Beta distribution that determines the cell type of the neurite. The maximum likelihood procedure for estimating α and β yields the following set of coupled equations:

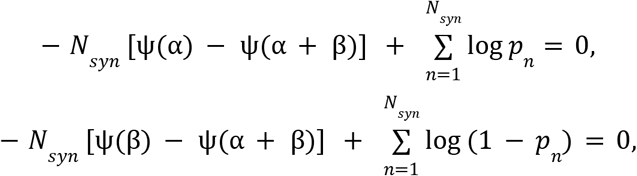

where ψ(.) is the digamma function. We can easily establish the following identity for the expected log odds of the excitatory class, 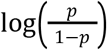, using the basic properties of Beta distribution:

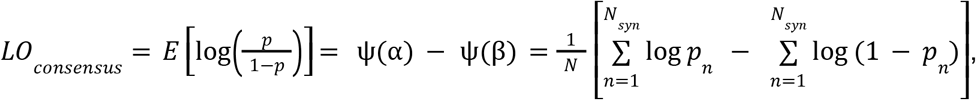

where the last equality results from maximum likelihood equations given earlier. According to this simple model, the consensus procedure amounts to summing up the log probabilities of excitatory class predictions and subtracting the log probabilities of inhibitory class predictions. We make a consensus cell type call for a neurite depending on the expected log odds: inhibitory if *LO*_*consensus*_ > 0, and excitatory if *LO*_*consensus*_ < 0.

#### Step 2 (Synapse Consensus Cell Typing)

We simply assign the consensus cell type of a neurite to the participating synapses according to the polarity of the synapse: if the neurite is the presynaptic partner of synapse, the neurite consensus cell type is thought of as the consensus presynaptic cell type of the synapse, and similarly for postsynaptic neurites.

### Software used

CloudVolume

https://github.com/seung-lab/cloud-volume

### Software referenced

MicronsBinder https://github.com/AllenInstitute/MicronsBinder

### Original software

SynapseCLR and SynapseAugmenter Python packages

Jupyter notebooks for recreating all of the figures in the paper

Code for visualizing image chunks and segmentation layers in Python

Code for assigning consensus cell types to neurites and synapses

Code for imputation and prediction cross-validation of synapse annotations using SynapseCLR representations

https://github.com/broadinstitute/SynapseCLR

## SUPPLEMENTARY FIGURES

**Figure S1.**
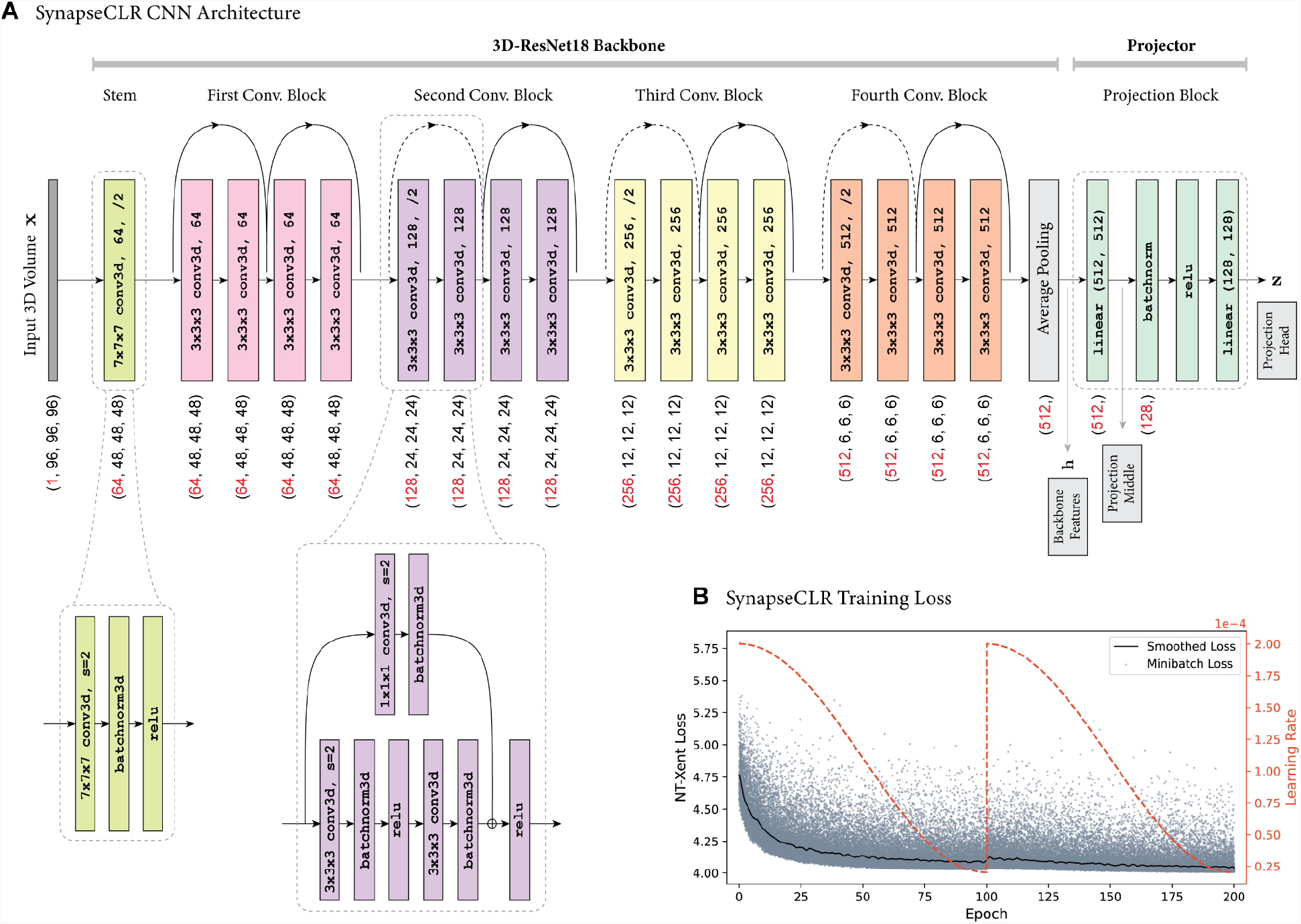
SynapseCLR network architecture and training loss. (A) The architecture of the 3D-ResNet18 with CIFAR stem and four convolutional blocks with residual connection. The CIFAR stem is a 7×7×7 3D convolution with stride 2, followed by batch normalization and ReLU activation (shown below the main network). Dotted shortcut lines correspond to downsampling via stride-2 convolutions before element-wise residual addition prior to activation (also shown below the main network). Black shortcut lines are similar to dotted shortcuts, though, without the downsampling operation. The projection block we use is a 2-layer perceptron with batch normalization and ReLU activation. Backbone (***h***), projection middle, and projection head (***z***) features are indicated. (B) The SynapseCLR NT-Xent training loss per minibatch is shown in gray (the smoothed loss is shown in black). We train the network for 200 epochs with cosine learning rate decay and a restart after 100 epochs (Methods).

**Figure S2.**
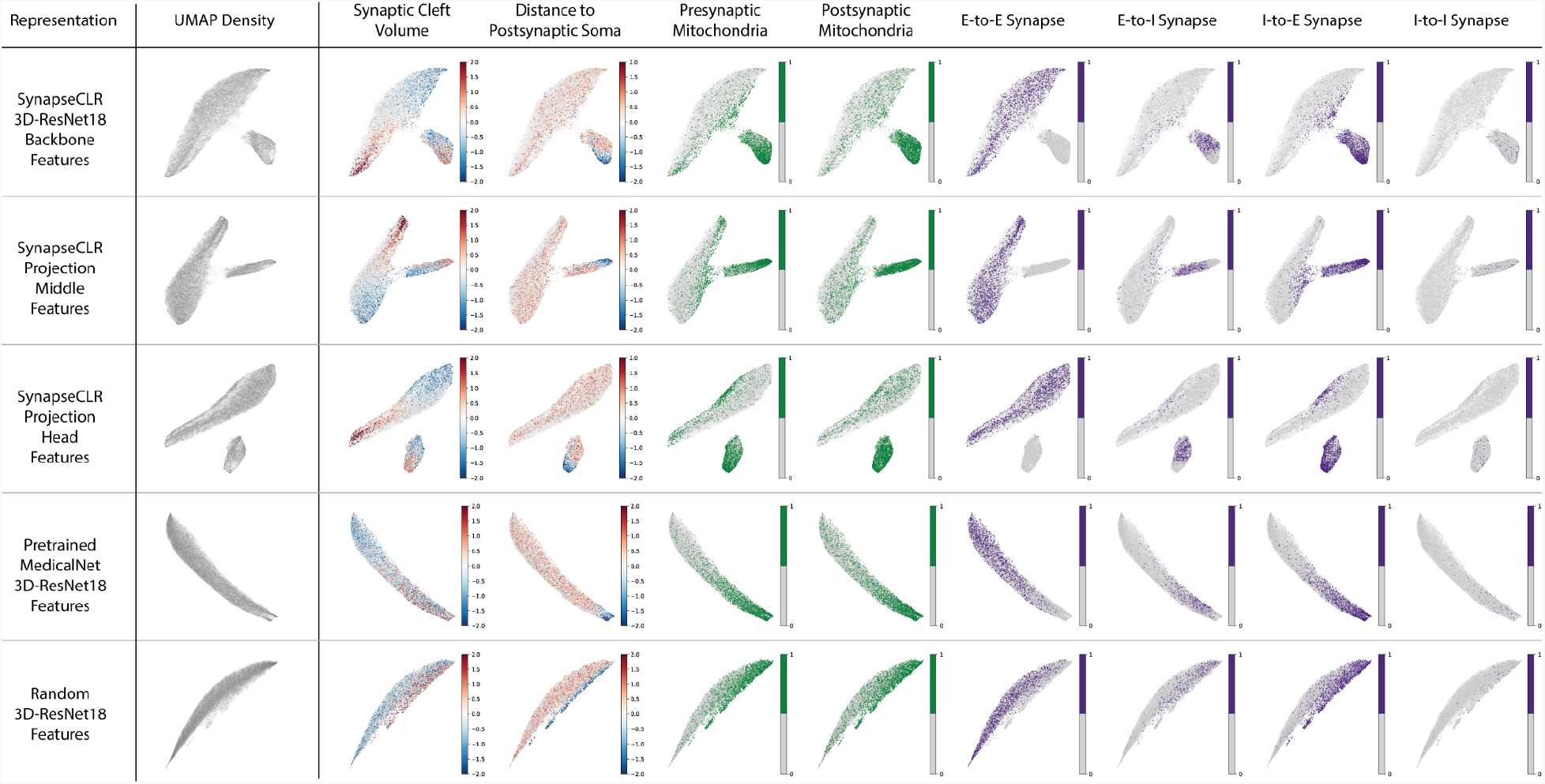
UMAP embeddings of different representations of synapse 3D EM data. The first three rows correspond to representations extracted from the 3D-ResNet18 backbone, projection middle, and projection head of SynapseCLR network (see Fig. S1A). The last two rows correspond to features extracted from a pre-trained general-purpose 3D-ResNet18 (MedicalNet), and an untrained 3D-ResNet18 (random convolutional features). The columns show the UMAP embedding density, followed by several colorations according to the annotations available for a subset of synapses. Note the structural similarity of all three SynapseCLR representations, and the relative absence of structure in the two shown baselines.

**Figure S3.**
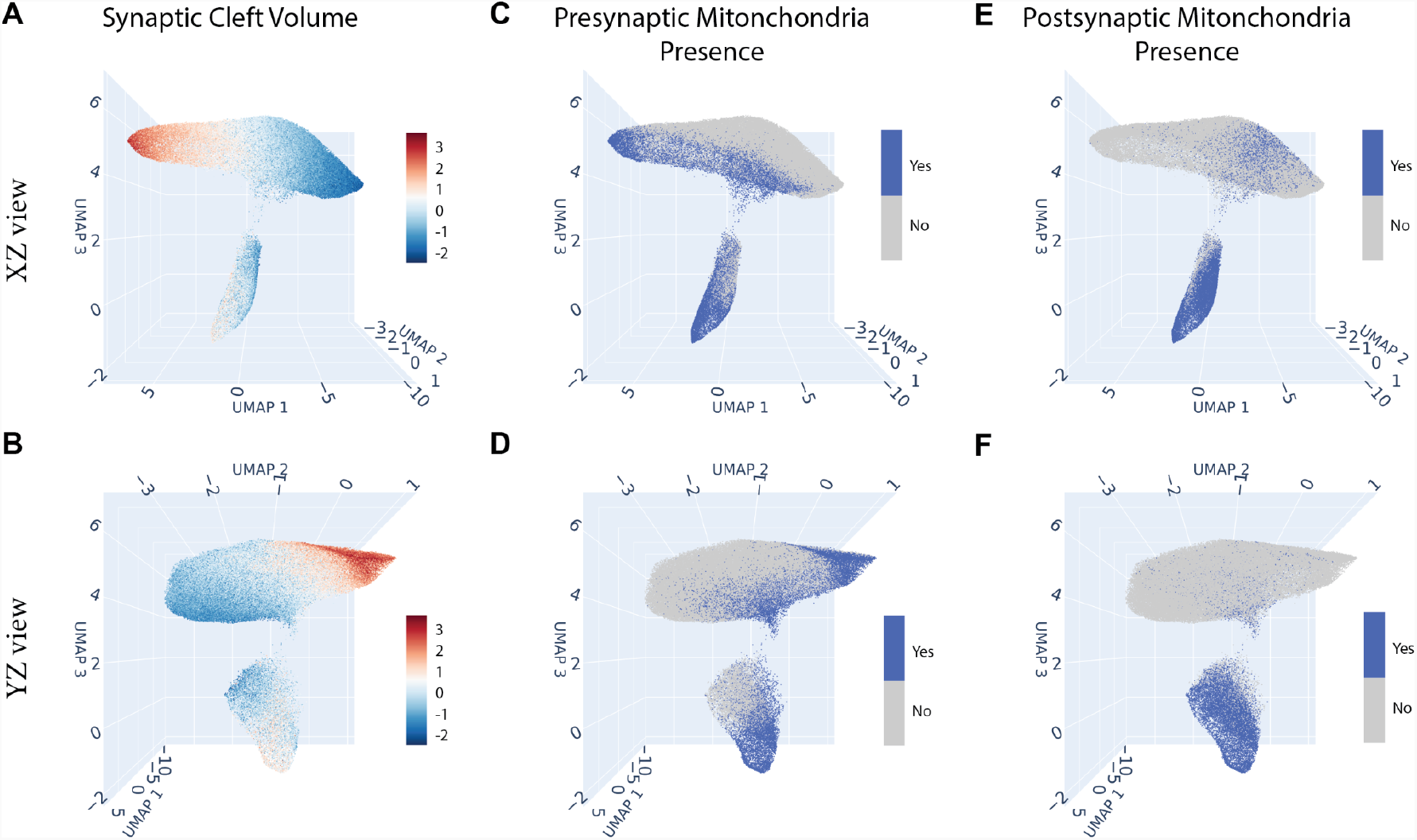
3D UMAP embedding of SynapseCLR backbone representations colored by three different annotations. The shown annotations are imputed using Gaussian process (GP) regression (Methods). Each column corresponds to a different annotation. For each annotation, the top and bottom rows show viewpoints aimed at UMAP1-UMAP3 and UMAP2-UMAP3 planes, respectively.

**Figure S4.**
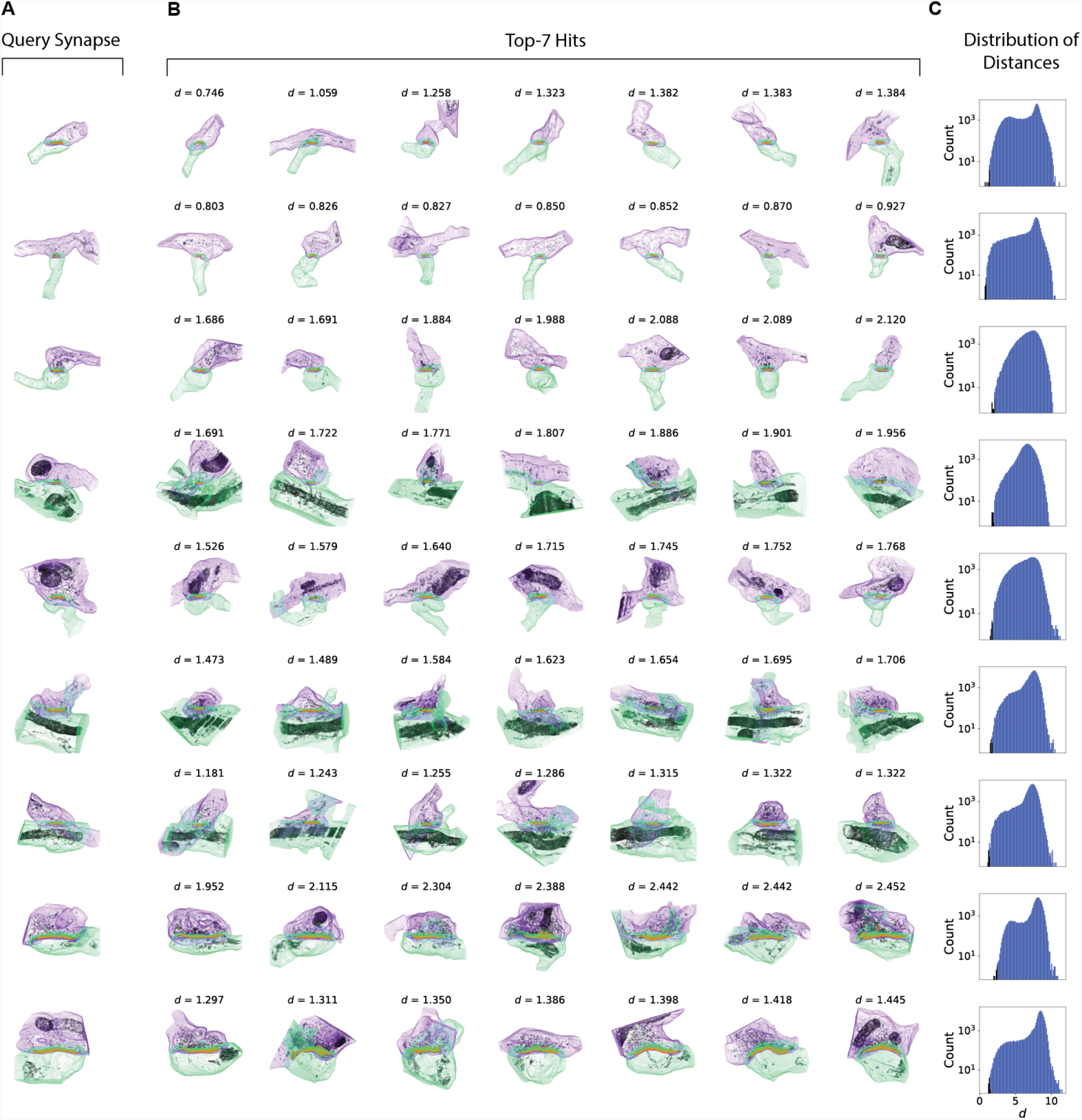
Additional examples of distance-based similarity queries from the SynapseCLR representation space (see Fig. 2D-F). (A) The “prompt” synapses for our queries. (B) Top-7 queried synapses, corresponding to the top-7 nearest synapses according to Euclidean distance in the SynapseCLR representation space. (C) The histogram of Euclidean distances between the prompt synapse and all other synapses in our dataset. The top-7 synapses shown in (B) are highlighted in black in the histogram.

**Figure S5.**
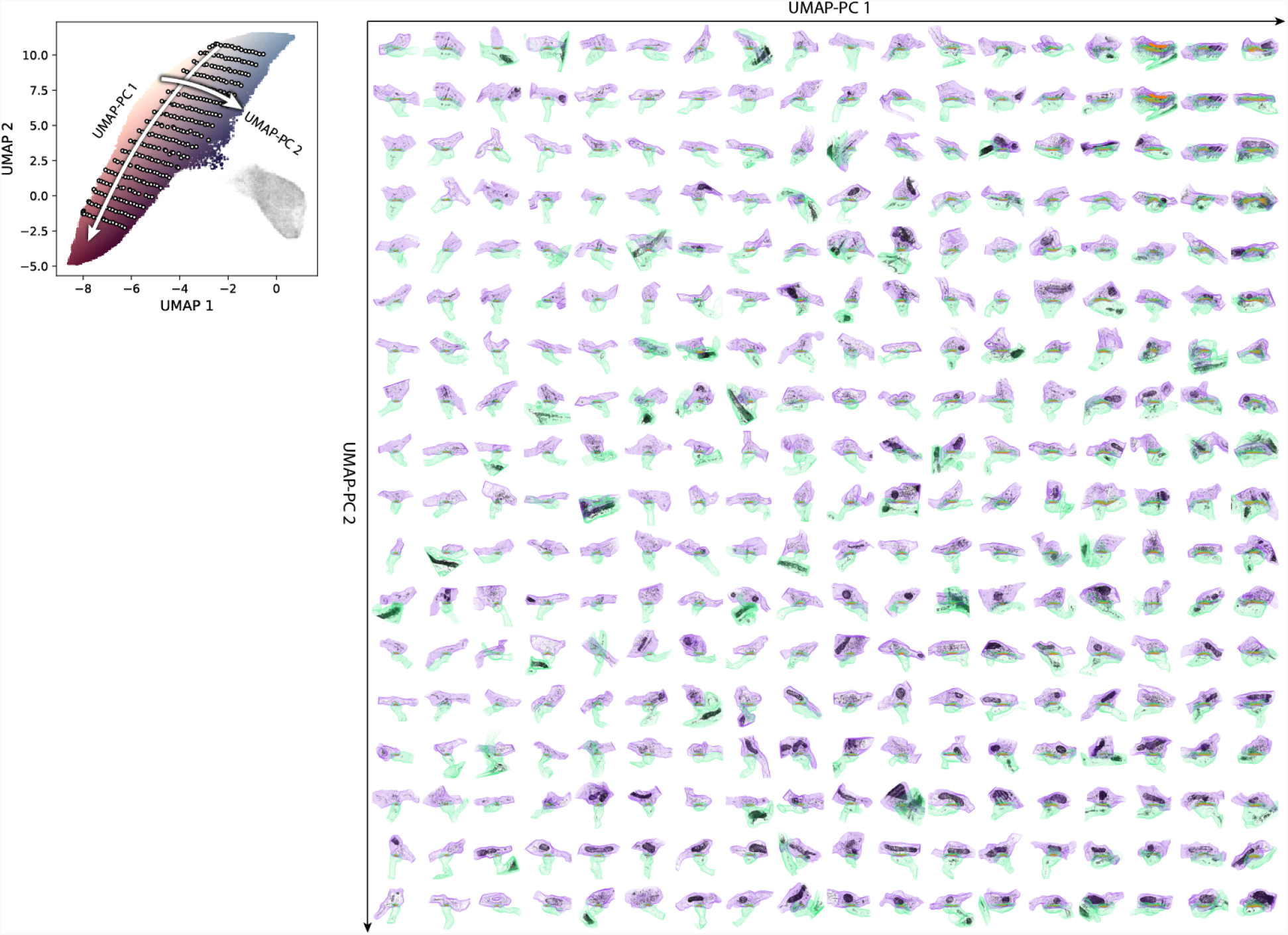
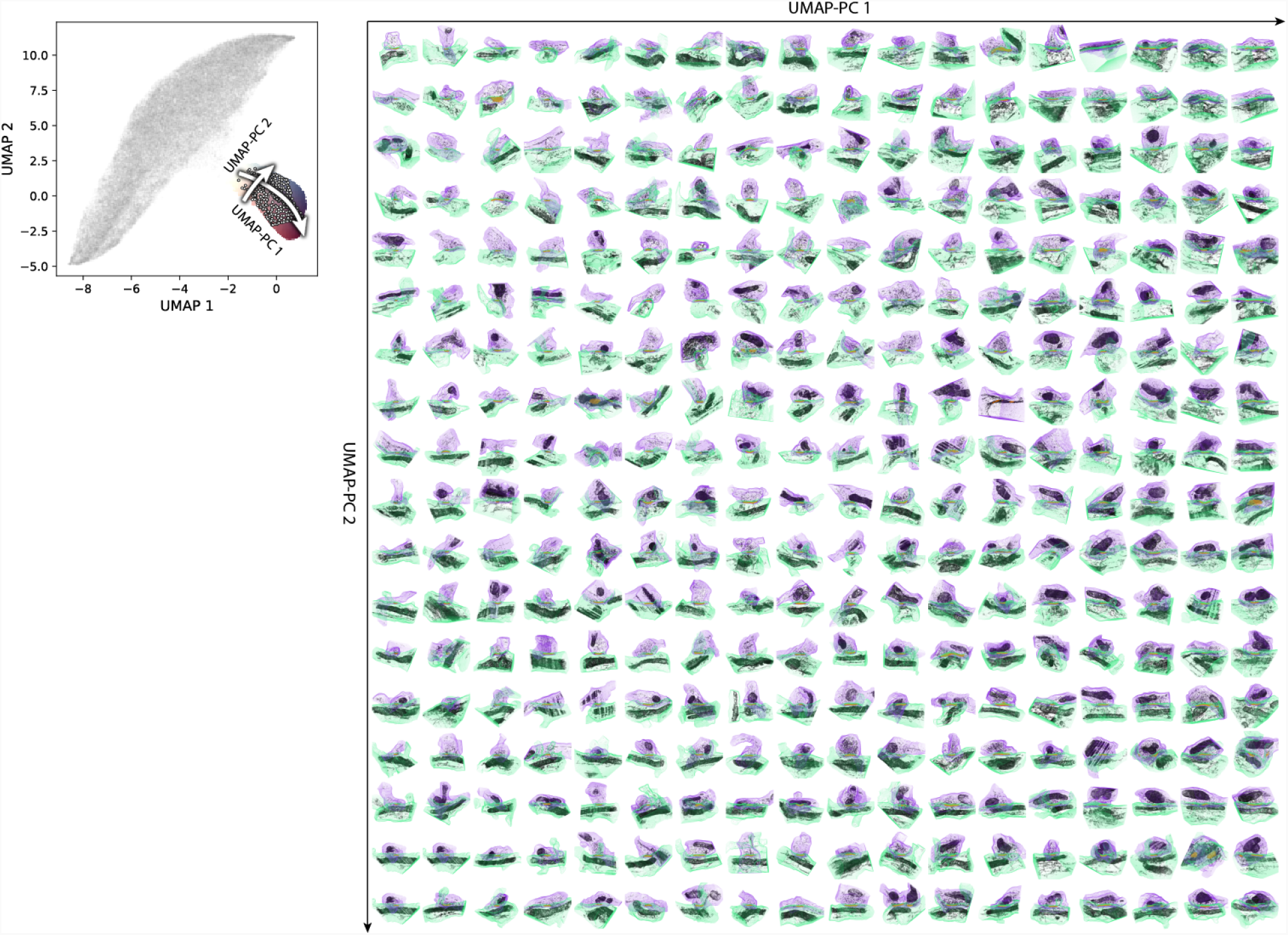
Synapse illustrations over a finer two-dimensional UMAP-PC grid over the excitatory sub-manifold (see Fig. 5B). Synapse illustrations over a finer two-dimensional UMAP-PC grid over the inhibitory sub-manifold (see Fig. 5F).

**Figure S6.**
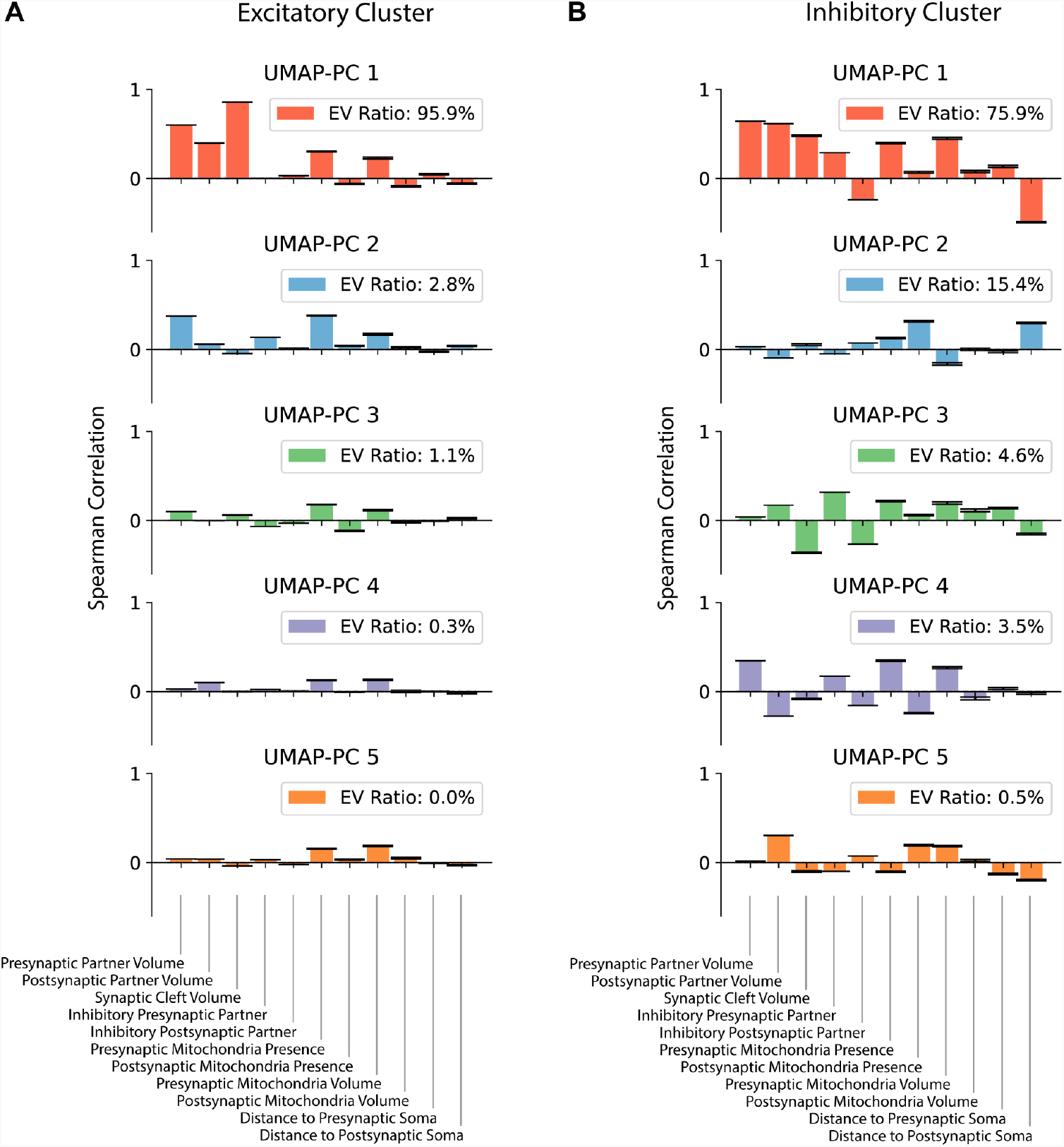
Five-dimensional UMAP-PCA analysis of the excitatory and inhibitory UMAP sub-manifolds of SynapseCLR backbone representations. Explained variance (EV) ratio of each of the UMAP-PCs is shown in the legend, along with the Spearman correlation with each of the available annotations. UMAP-PCs in the 3rd and higher dimensions explain a negligible amount of variance. (A) UMAP-PCA analysis of the excitatory sub-manifold. (B) UMAP-PCA analysis of the inhibitory sub-manifold.

**Figure S7.**
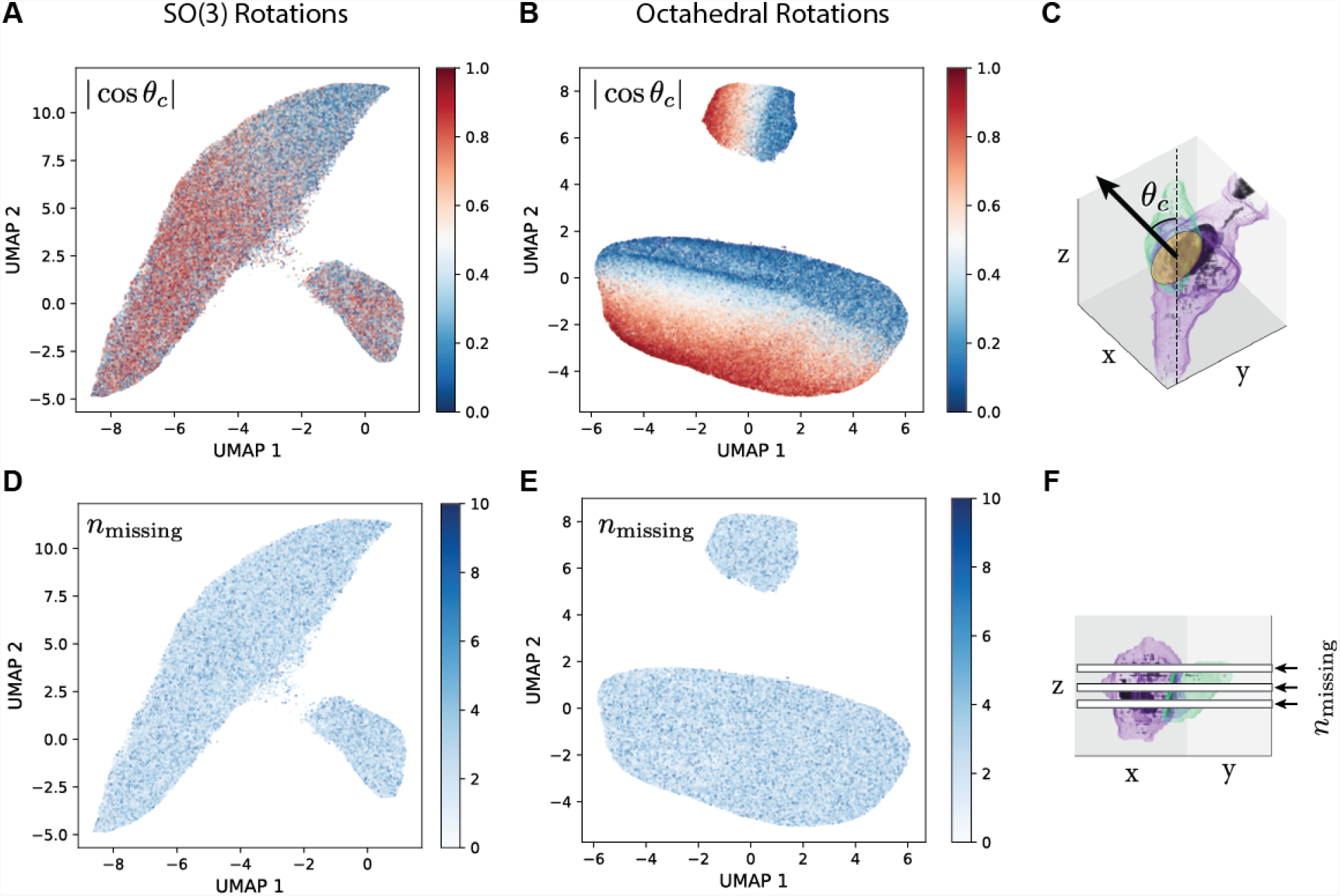
The effect of insufficient data augmentation on SynapseCLR representations. Panels (A) and (B) show the UMAP embeddings of SynapseCLR backbone features colored by the cosine of the angle between the synaptic cleft plane normal vector and the z-axis, as illustrated in panel (C). Representations obtained by allowing arbitrary SO(3) rotations are shown in (A) and are largely insensitive to the orientation of the synaptic cleft. In contrast, (B) shows representations obtained by allowing only discrete octahedral rotations and reflections. The short axis of variation in each sub-manifold is entirely driven by the synaptic orientation, which is an undesirable and, in general, a biologically irrelevant feature. Panels (D) and (E) demonstrate the efficacy of augmentation as a strategy in general in both training runs: they show the same UMAPs as (A) and (B), but colored based on the number of missing axial planes in each synapse image, as illustrated in panel (F). In both runs, we included a data augmentation mimicking this artifact and as a result, both representations are insensitive to the number of blank (missing) z-sections.

**Figure S8.**
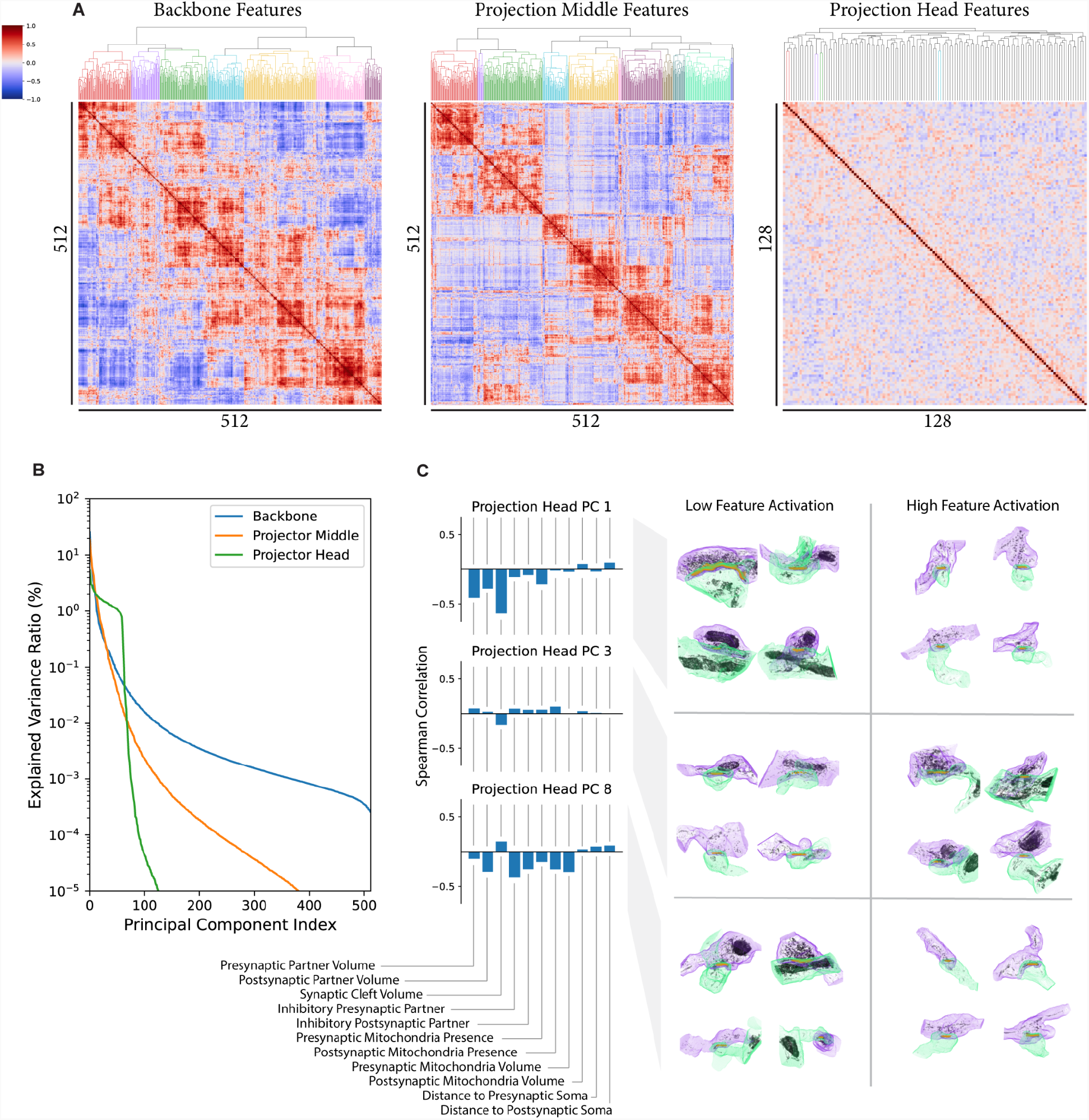
The correlation structure of various SynapseCLR features and the interpretability of projection head features. (A) Clustered correlation heatmaps for the SynapseCLR backbone (left), projection middle (middle), and projection head (right) features (see Fig. S1). Note that features are progressively disentangled as they are processed through the projector, with the projection head showing nearly completely independent features. (B) PCA of SynapseCLR backbone, projection middle, and projection head features, showing that approximately 60 out of 128 projection head features are linearly independent. (C) Many projection head PCs have emergent structural interpretations: lower values of PC 1 in a synapse correspond to larger synapses and containment of mitochondria; higher values of PC 3 seemingly correspond to the presence of presynaptic vesicles, a measurement that is not part of our manual annotations; finally, lower values of PC 8 correspond to presence of pre- and postsynaptic mitochondria and their volumes, with significantly reduced dependence on synaptic cleft and partner volumes compared to PC 1.

## SUPPLEMENTARY TABLES

**Table S1.** The effect of hyperparameters and choice of SynapseCLR features on the cross-validation accuracy of Gaussian process (GP) regression for the imputation of synapse annotations. The first group of rows (3 in total) corresponds to using different SynapseCLR features. The next group of rows (3 in total) shows the effect of PCA dimensionality reduction of features prior to performing regression. Finally, the last group of rows (22 in total) shows the effect of different kernel functions and different numbers of inducing points. The top-performing model for each synaptic annotation is highlighted in bold within each of the three row groupings.

| Kernel Type | PCA | Feature | Number of Inducing Points | Synaptic Cleft Volume (EV) | Distance from Presynaptic Soma (EV) | Distance from Postsynaptic Soma (EV) | Presynaptic Mitochondria Volume (EV) | Postsynaptic Mitochondria Volume (EV) | Presynaptic Cell Type (AUC) | Postsynaptic Cell Type (AUC) | Presynaptic Mitochondria Presence (AUC) | Postsynaptic Mitochondria Presence (AUC) |
| --- | --- | --- | --- | --- | --- | --- | --- | --- | --- | --- | --- | --- |
| RBF | No | Projection Head | 50 | 60.964 ± 3.280 | 8.246 ± 1.905 | 49.055 ± 0.810 | 48.776 ± 4.049 | 2.821 ± 2.113 | 0.954 ± 0.009 | 0.918 ± 0.005 | 0.914 ± 0.023 | 0.829 ± 0.004 |
| RBF | No | Projection Middle | 50 | 66.056 ± 2.935 | 13.675 ± 1.931 | 51.367 ± 1.125 | 53.930 ± 4.064 | 5.943 ± 2.812 | 0.966 ± 0.005 | 0.931 ± 0.002 | 0.931 ± 0.014 | 0.833 ± 0.014 |
| RBF | No | Backbone | 50 | **68.779 ± 2.871** | **15.812 ± 1.549** | **53.551 ± 1.497** | **56.184 ± 3.349** | **8.733 ± 2.801** | **0.974 ± 0.004** | **0.939 ± 0.001** | **0.936 ± 0.016** | **0.851 ± 0.011** |
| RBF | 50 | Backbone | 50 | 65.534 ± 3.117 | 11.983 ± 2.277 | 51.576 ± 1.445 | 52.847 ± 3.722 | 6.867 ± 2.049 | 0.967 ± 0.003 | 0.929 ± 0.003 | 0.931 ± 0.014 | 0.839 ± 0.009 |
| RBF | 100 | Backbone | 50 | 66.575 ± 3.125 | 13.979 ± 1.659 | 52.476 ± 1.375 | 54.113 ± 4.136 | 7.674 ± 2.346 | 0.972 ± 0.003 | 0.934 ± 0.003 | 0.935 ± 0.015 | 0.847 ± 0.011 |
| RBF | No | Backbone | 50 | **68.779 ± 2.871** | **15.812 ± 1.549** | **53.551 ± 1.497** | **56.184 ± 3.349** | **8.733 ± 2.801** | **0.974 ± 0.004** | **0.939 ± 0.001** | **0.936 ± 0.016** | **0.851 ± 0.011** |
| Linear | No | Backbone | 5 | 65.684 ± 3.187 | 14.512 ± 2.205 | 51.837 ± 1.215 | 51.904 ± 3.955 | 9.525 ± 2.281 | 0.957 ± 0.006 | 0.914 ± 0.003 | 0.922 ± 0.019 | 0.836 ± 0.019 |
| Linear | No | Backbone | 10 | 66.428 ± 3.102 | 14.740 ± 2.054 | 52.391 ± 1.172 | 53.147 ± 3.906 | 9.852 ± 2.473 | 0.965 ± 0.005 | 0.924 ± 0.004 | 0.928 ± 0.017 | 0.839 ± 0.017 |
| Laplace | No | Backbone | 10 | 64.708 ± 3.190 | 12.313 ± 1.795 | 50.875 ± 0.536 | 50.370 ± 4.085 | 9.470 ± 1.828 | 0.955 ± 0.007 | 0.914 ± 0.003 | 0.921 ± 0.019 | 0.830 ± 0.013 |
| Laplace | No | Backbone | 20 | 65.784 ± 3.272 | 13.857 ± 1.815 | 51.687 ± 0.893 | 51.977 ± 4.144 | 9.253 ± 1.843 | 0.963 ± 0.006 | 0.923 ± 0.005 | 0.924 ± 0.019 | 0.834 ± 0.013 |
| Laplace | No | Backbone | 50 | 66.458 ± 3.193 | 14.713 ± 2.007 | 52.000 ± 0.858 | 53.222 ± 3.771 | 9.109 ± 2.095 | 0.965 ± 0.006 | 0.928 ± 0.003 | 0.926 ± 0.018 | 0.837 ± 0.012 |
| Laplace | No | Backbone | 100 | 66.678 ± 3.197 | 14.365 ± 1.741 | 52.064 ± 1.107 | 53.727 ± 3.622 | 8.832 ± 2.115 | 0.966 ± 0.005 | 0.932 ± 0.003 | 0.927 ± 0.017 | 0.839 ± 0.011 |
| Laplace | No | Backbone | 200 | 66.735 ± 3.139 | 14.059 ± 1.662 | 52.333 ± 0.932 | 53.787 ± 3.524 | 7.840 ± 2.350 | 0.967 ± 0.005 | 0.933 ± 0.003 | 0.928 ± 0.017 | 0.839 ± 0.012 |
| Laplace | No | Backbone | 500 | 66.935 ± 3.016 | 13.800 ± 1.768 | 52.343 ± 0.944 | 54.098 ± 3.307 | 6.165 ± 2.787 | 0.968 ± 0.005 | 0.935 ± 0.005 | 0.929 ± 0.018 | 0.842 ± 0.011 |
| RBF | No | Backbone | 5 | 64.984 ± 3.287 | 13.588 ± 1.876 | 51.616 ± 1.202 | 51.885 ± 3.941 | 9.556 ± 2.048 | 0.965 ± 0.005 | 0.922 ± 0.005 | 0.897 ± 0.059 | 0.838 ± 0.016 |
| RBF | No | Backbone | 10 | 66.533 ± 3.104 | 14.392 ± 1.872 | 52.224 ± 1.144 | 53.350 ± 3.824 | **9.968 ± 2.402** | 0.969 ± 0.005 | 0.930 ± 0.005 | 0.931 ± 0.016 | 0.841 ± 0.014 |
| RBF | No | Backbone | 20 | 67.531 ± 2.956 | 15.284 ± 1.803 | 52.763 ± 1.176 | 54.531 ± 3.775 | 9.804 ± 2.568 | 0.973 ± 0.004 | 0.938 ± 0.002 | 0.933 ± 0.016 | 0.847 ± 0.014 |
| RBF | No | Backbone | 50 | 68.779 ± 2.871 | 15.812 ± 1.549 | 53.551 ± 1.497 | 56.184 ± 3.349 | 8.733 ± 2.801 | 0.974 ± 0.004 | 0.939 ± 0.001 | 0.936 ± 0.016 | 0.851 ± 0.011 |
| RBF | No | Backbone | 100 | 69.454 ± 2.791 | **15.938 ± 1.426** | 53.882 ± 1.580 | 56.489 ± 3.138 | 7.398 ± 2.515 | **0.975 ± 0.004** | **0.940 ± 0.004** | 0.938 ± 0.016 | 0.854 ± 0.010 |
| RBF | No | Backbone | 200 | 69.882 ± 2.683 | 15.481 ± 1.244 | **53.902 ± 1.584** | 56.785 ± 3.144 | 5.441 ± 2.534 | 0.975 ± 0.004 | 0.940 ± 0.005 | **0.939 ± 0.015** | 0.856 ± 0.010 |
| RBF | No | Backbone | 300 | 69.966 ± 2.637 | 14.768 ± 1.023 | 53.801 ± 1.745 | **56.869 ± 3.091** | 4.002 ± 2.583 | 0.975 ± 0.003 | 0.939 ± 0.006 | 0.939 ± 0.015 | **0.857 ± 0.010** |
| RBF | No | Backbone | 400 | **70.017 ± 2.669** | 14.371 ± 0.969 | 53.791 ± 1.853 | 56.824 ± 3.013 | 2.770 ± 2.611 | 0.975 ± 0.004 | 0.938 ± 0.007 | 0.939 ± 0.015 | 0.856 ± 0.011 |
| RBF | No | Backbone | 500 | 70.003 ± 2.635 | 13.930 ± 0.794 | 53.645 ± 2.002 | 56.861 ± 2.975 | 1.990 ± 2.868 | 0.975 ± 0.004 | 0.937 ± 0.008 | 0.939 ± 0.015 | 0.855 ± 0.011 |
| RBF | No | Backbone | 600 | 69.950 ± 2.622 | 13.597 ± 0.797 | 53.612 ± 2.048 | 56.759 ± 2.926 | 1.042 ± 2.806 | 0.975 ± 0.004 | 0.936 ± 0.008 | 0.938 ± 0.015 | 0.855 ± 0.011 |
| RBF | No | Backbone | 700 | 69.921 ± 2.587 | 13.552 ± 0.693 | 53.588 ± 2.004 | 56.786 ± 2.992 | -0.243 ± 3.136 | 0.975 ± 0.004 | 0.936 ± 0.009 | 0.938 ± 0.015 | 0.854 ± 0.011 |
| RBF | No | Backbone | 800 | 69.954 ± 2.542 | 13.230 ± 0.529 | 53.400 ± 2.052 | 56.537 ± 2.979 | -0.926 ± 2.916 | 0.975 ± 0.004 | 0.935 ± 0.009 | 0.938 ± 0.015 | 0.853 ± 0.011 |
| RBF | No | Backbone | 900 | 69.905 ± 2.598 | 12.860 ± 0.734 | 53.297 ± 2.155 | 56.473 ± 2.928 | -1.259 ± 3.019 | 0.975 ± 0.004 | 0.935 ± 0.009 | 0.939 ± 0.016 | 0.851 ± 0.011 |
| RBF | No | Backbone | 1000 | 69.908 ± 2.554 | 12.603 ± 0.568 | 53.197 ± 2.103 | 56.489 ± 2.895 | -1.427 ± 3.233 | 0.975 ± 0.004 | 0.934 ± 0.009 | 0.938 ± 0.016 | 0.850 ± 0.011 |

